# Multimodal imaging reveals multiscale mechanical interplay in vertebral endplate microarchitecture during intervertebral disc loading

**DOI:** 10.1101/2024.08.19.608559

**Authors:** Alissa L. Parmenter, Elis Newham, Aikta Sharma, Catherine M. Disney, Hans Deyhle, Federico Bosi, Nick J. Terrill, Brian K. Bay, Andrew A. Pitsillides, Himadri S. Gupta, Peter D. Lee

## Abstract

The function of all musculoskeletal joints depends on hierarchical structures spanning the molecular to whole joint scales. Investigating biomechanics across length scales requires correlative multiscale experimental methods. This study applies multimodal *in situ* synchrotron imaging techniques to spinal joints – focussing on the vertebral endplates – to explore relationships between structure and mechanical strain across spatial scales. Strain mapping using digital volume correlation combined with microarchitectural analysis reveals that high tensile and shear strains play a role in the cartilage to bone transition. Correlative imaging and diffraction show that bone contains narrower mineral nano-crystallites under greater compressive prestrain compared to calcified cartilage. We hypothesise that this multiscale structural adaptation supports the mechanical function of the intervertebral disc. Future applications of the techniques presented here have potential to help unravel biomechanical underpinnings of pathologies affecting mineralised tissue structure. The multiscale structure-function relationships uncovered here may inspire the design of biomaterials and orthopaedic implants.

## INTRODUCTION

The multiscale structure of bone and cartilage underpins the biomechanical function of the musculoskeletal system. In the spine, bone and cartilage at the vertebral surface – the vertebral endplates (VEPs) – help control load and nutrient transfer from the intervertebral discs (IVDs). Back pain affects 600 million people worldwide and is the leading cause of years lived with disability (due to increase by 36 % by 2050) ^1^. Spine biomechanics plays a critical role in back pain related pathologies such as intervertebral disc (IVD) degeneration^2^ and scoliosis^3^. There is therefore great interest in the biomechanics of the spine and its constituent parts, including the IVDs and vertebral endplates (VEPs)^4^, which are highly complex across hierarchical spatial scales (Fig. 1) and experience a large range of movements and mechanical loads in day-to-day life.

**Figure 1.**
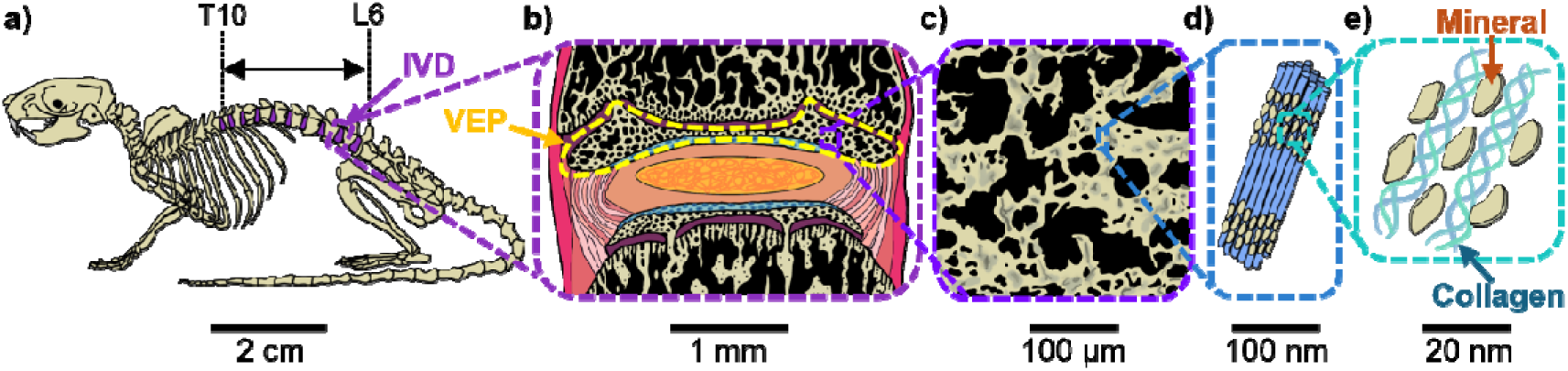
Structural hierarchy in the murine vertebral endplates. Illustrations of a) the murine skeleton with IVDs investigated in this study, from thoracic vertebra 10 (T10) to lumbar vertebra 6 (L6), shown in purple; b) the murine intervertebral disc, with cranial vertebral endplate outlined in yellow; c) calcified tissue microstructure; d) a mineralised collagen fibril; e) collagen fibres and mineral platelets

The IVD, comprising of the collagenous annulus fibrosus (AF) and proteoglycan rich nucleus pulposus (NP), deforms under loading, applying complex, location specific strains to the calcified VEPs (Fig. 1b). The IVD receives nutrition through canals in the VEPs^5^, therefore VEP structure must balance mechanical competency with permeability. The VEPs are comprised of calcified cartilage and bone; these mineralised tissues have differing material properties, with the elastic modulus of calcified cartilage an order of magnitude lower than bone^6^. The presence of a calcified cartilage layer between the soft tissues of the IVD and stiff vertebral bone helps create a smoother transition in material properties, reducing high strain concentrations at tissue boundaries. This greater bone stiffness persists even when mineral content is matched to calcified cartilage levels^7^, suggesting that inherent differences in non-mineralised components influences their respective mechanical properties. Raman spectroscopy has shown that there are differences in both the organic and mineral composition in calcified cartilage compared to subchondral bone in human osteochondral junctions, including higher intensity glycosaminoglycan, lipid, phosphate and carbonate peaks in calcified cartilage^8^. Location and structure of the mineralised crystallites within the collagenous network^9–17^ will play a crucial role in the local mechanical equilibrium and load transfer between the mineral and collagen, and we hypothesise that this differs between calcified cartilage and bone because of inherent differences in their organic components.

The murine VEP forms a calcified cartilage epiphysis with distinct secondary ossification (bone formation) centres, separated from the adjacent vertebral bone by a cartilaginous growth plate^18^. The formation of different musculoskeletal tissues is partly driven by the local mechanical environment^19,20^. Osteocytes in the bone compartment are considered responsive to prevailing mechanical cues and influence the three-dimensional structure of the VEPs ^21,22^. Chondrocytes in cartilage are also mechanosensitive, with changes in their mechanical environment known to lead to degeneration^23^. VEP injury and degeneration disrupts the mechanical environment, driving IVD degeneration and back pain^24,25^, whilst VEP maintenance in health relies on the formation of optimal structures for load absorption and transfer. An appreciation of how these structures attain their function requires investigation of their biomechanics during the growth stage.

The mechanical function of VEPs is impacted by structures across spatial scales. At the macroscale, VEP thickness influences their response to load (Fig. 1b). At the microscale, mineralised tissue architecture and tissue mineral density (TMD, the amount of mineral in a given volume of mineralised tissue) dominate (Fig. 1c), with microstructure linked to vertebral failure ^26^. At the nanoscale, mineralised collagen fibril structure (Fig. 1d-e) and prestrain impact mechanical properties^17,27^. Prestrain - the inherent strain in a material in its undeformed state - plays a vital mechanical role ^2,28,29^; loss of prestrain in the AF leads to IVD degeneration ^2^. Mineralised tissue prestrain is thought to increase toughness and crack resistance^30–32^, and is partly caused by the contraction of collagen fibrils during the biomineralization process^33,34^. Cells have been shown to actively generate tension in the extracellular matrix^20,35^, and this may confer prestrain in the mineral. Mineral crystallites make up 40-45% of the bone matrix volume, with the degree and nature of collagen mineralisation leading to large changes in tissue mechanical properties^27^. Mineral structure across the VEPs remains unknown and we predict that this structure influences whole joint level biomechanics.

VEP and IVD physiology thus involves a complex feedback system between structure and mechanical forces across a range of length scales. We hypothesise that a combination of VEP microarchitecture, mineral nanostructure, and prestrain enables optimal IVD mechanical function and is partly controlled by load induced strains *in vivo*.

Several techniques can be used to characterise biomechanics at particular spatial scales - on the whole joint to microscale, synchrotron X-ray computed tomography (sCT) and digital volume correlation (DVC) are powerful tools for investigating the strain response of musculoskeletal tissues ^36–41^, and have been used to measure microscale strains in trabecular bone^39,40^. However, while DVC has been applied to study internal IVD ^42,43^ and vertebral bone ^44^ strains, microscale strain patterns in the VEPs have not been examined. The analysis of microscale strain fields across whole joints using DVC requires imaging with both a high spatial resolution and a large field of view. At the nanoscale, wide-angle X-ray diffraction (WAXD) can be used to quantify mineral prestrain^17^ and crystallite size^14^, and has been used to investigate fracture toughness in bone^45^ and dentin^30^, as well as how mineral crystal structure changes with osteoarthritis^15^. 2D WAXD gives a diffraction signal with contributions from everything along the X-ray beam path, making interpretation difficult for complex, heterogeneous sample geometries. 3D WAXD methods help overcome this but require rotation of the sample around multiple axes and currently very long experimental times, making *in situ* experiments difficult^46^. There remains a need for hierarchical, multiscale imaging to bridge the gap between the cell mechanical environment and whole joint biomechanics that can reveal the relationships between mineralised tissue prestrain and whole joint level mechanics. Microscale load-induced strains in the VEP during whole IVD loading and how these strains are related to microarchitecture remain unknown, with previous mechanical studies on VEPs involving destructive sample preparation^47–49^ or imaging methods unable to resolve 3D microscale features^50–53^. Variation in mineral structure and prestrain across the VEPs has not previously been measured, and the direct correlation between nanoscale mineral properties and the microarchitecture of mineralised tissues is difficult to observe.

This study aims to uncover relationships between VEP structure and mechanical strain from the nano- to whole-joint-scales using correlative synchrotron imaging techniques, demonstrating a novel combination of methodologies for evaluating structure-strain relationships across spatial scales. We present a series of methodological improvements for imaging whole IVD *in situ* and perform correlative sCT and WAXD of VEPs at the Dual Imaging and Diffraction (DIAD) beamline^54^ of the Diamond Light Source (DLS) synchrotron. By combining the analysis of these two experiments, multiscale structure-function relationships can be uncovered.

We use sCT and DVC analysis to investigate whole joint to microscale structure and mechanics in intact murine IVDs under physiologically relevant loading. We measure relative TMD, microarchitecture, and nanoscale-precision local load-induced strains to evaluate structure-strain relationships in whole VEPs. Microarchitectural values are used to determine predominant tissue type in each region, enabling comparisons between calcified cartilage and bone without complex segmentation. Micro-to-nanoscale (Fig. 1c-e) structure and prestrain are measured by spatially correlated sCT and WAXD, to unveil relationships between microarchitecture, relative TMD, mineral structure and prestrain. Because mineral particles are elastic and much stiffer than the organic phase (mineral Youngs Modulus ∼100 GPa vs ∼1 - 2 GPa for collagen), the stress on the mineral is proportional to the change in lattice spacing (d-period)^17^. Thus, relative changes in d-period are proportional to the mineral prestrain state and are larger in regions of tensile stress and smaller in regions of compressive stress. Hence, by measuring relative changes in d-period and crystallite structure across the osteochondral junction in the VEP, we will uncover local nanoscale mechanics. These studies yield novel insights in vertebral biology and mineralised tissue biomechanics.

## RESULTS

### sCT resolves microscale cell-matrix organisation and nanoscale precision strains by DVC

To achieve microscale resolution of cell-matrix organisation in whole IVDs under physiologically relevant loading conditions, phase contrast enhanced sCT scans of 9 murine spine segments under uniaxial compressive load were performed at beamline I13-2 at the DLS synchrotron (Fig. 2a,b). This built on previous work ^40,42,55,56^ (Methods), optimising experimental parameters to improve image quality whilst reducing scan time and minimising radiation dose. By imaging intact, fresh-frozen, hydrated IVDs under axial compression, the normal function of IVD structures could be observed, whereby the nucleus pulposus (NP) applies hydrostatic pressure in all directions, placing fibres in the AF into tension and causing the endplates to bulge outwards^57^, generating compressive, tensile, and shear strains in the VEP (Fig 2c). Four sCT scans were conducted for each sample; the first under an initial compressive pre-strain of 1 N and the following three under cumulative displacements of 40 µm (Fig. 2d). A 1.6 µm voxel size and 4.2 mm x 3.5 mm field of view (FOV) enabled the microstructures of hard and soft tissues to be resolved across the entire IVD (Fig. 2f, g). This resulted in a spatial resolution of 5.5 µm, calculated from Fourier shell correlation analysis (Figure S3). This resolution of microscale features allowed the accurate topographical quantification of bone and calcified cartilage microarchitecture across the VEPs using commonly applied metrics (see Methods). Image greyscale value (GSV) was measured to investigate relative changes in TMD across the VEPs. Here, the VEPs are defined as the mineralised epiphysis between the vertebral growth plate and the IVD.

**Figure 2.**
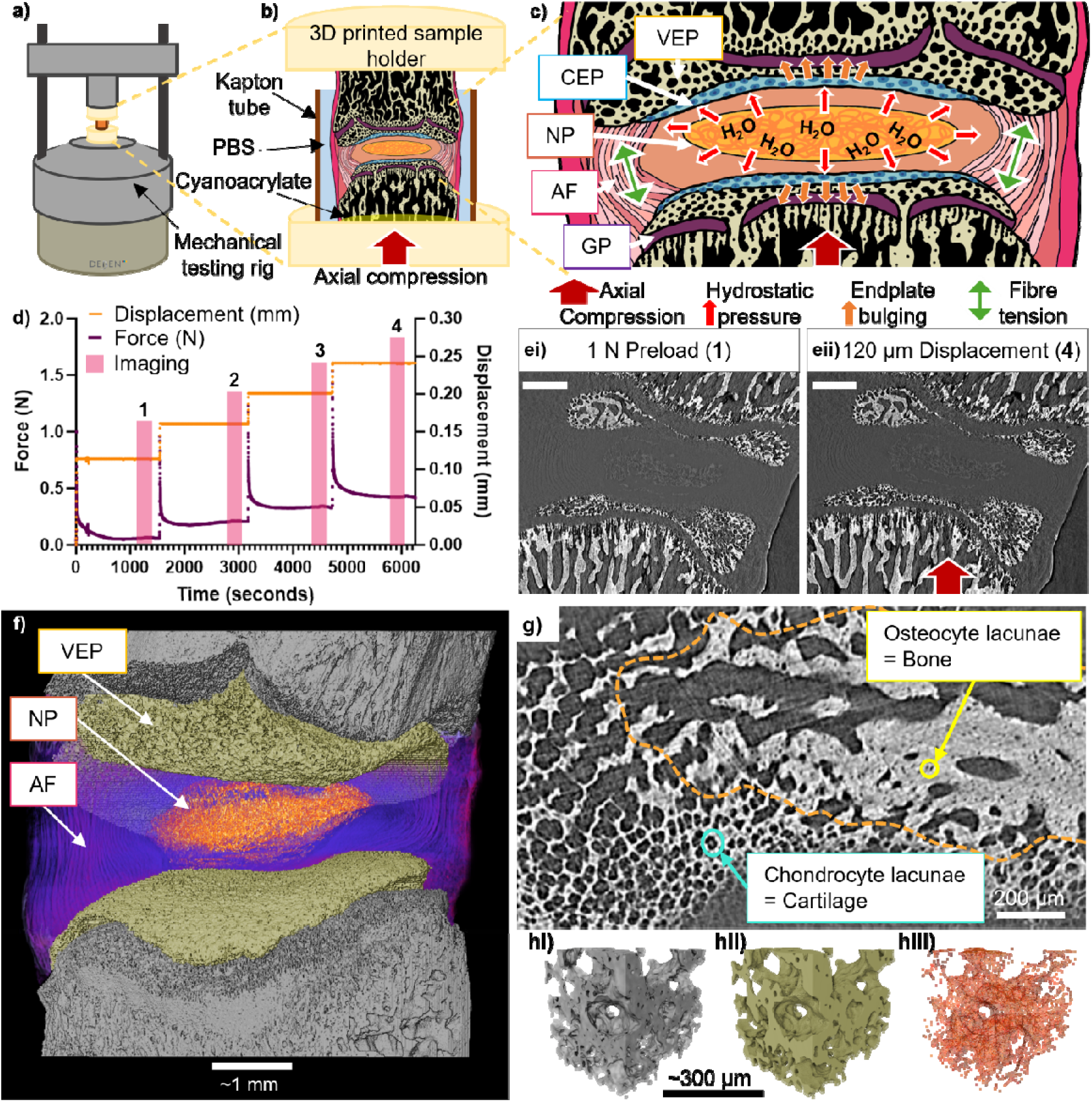
sCT resolves microscale cell-matrix organisation and nanoscale-precision strains by DVC. a) Schematic of the Deben CT500 mechanical testing rig and custom-built open frame with X-ray translucent carbon pillars to minimise imaging artefacts. b) Illustration of a cross-section of the sample environment, with 3D printed sample holder, Kapton tube, phosphate buffered saline (PBS), and cyanoacrylate glue labelled. c) Illustration of the IVDs response to axial compression, showing hydrostatic pressure in the nucleus pulposus (NP), fibre tension in the annulus fibrosus (AF), and bulging of the vertebral and cartilage endplates (VEP and CEP respectively) into the growth plate (GP). d) Force and displacement readings during one loading experiment, with readings taken during scanning highlighted in pink, scan 1 was taken 20 minutes after application of 1N preload, scan 2 after 4 % compression of the IVD, scan 3 after 8 % compression of the IVD, scan 4 after 12 % compression of the IVD. e) Sagittal image slice through a sample under i) 1 N preload, corresponding to scan 1 in d, and ii) 120 μm compression, corresponding to scan 4 in d, the direction of loading is shown by the red arrow, scalebars represent 0.5 mm. f) 3D rendering of a whole IVD sample, with the NP rendered in orange, AF in purple, VEPs in beige, and vertebral bone in grey. g) greyscale cross-section through a cranial VEP showing the microstructures of cartilage and bone, the bone compartment is approximately outlined in orange and identified by the presence of osteocyte lacunae, whereas the calcified cartilage contains chondrocyte lacunae. h) Point cloud generation for DVC, i) greyscale 3D rendering of VEP structure, ii) filtered and smoothed surface generated from VEP mineralised tissue structure, iii) tetrahedral grid of the surface, the nodes of the grid are used to define the centres of the subvolumes used for DVC image tracking.

To obtain 3D strain fields across the VEP, sCT datasets were subjected to DVC analysis^37,38^ using a local approach with microstructure specific point cloud (Figure 2h) and sub-voxel interpolation. This enabled the measurement of displacements with an accuracy of 10 – 85 nm and precision of 9 – 63 nm. Subsequent Lagrangian strains were calculated from the displacement field with an accuracy of 0.036 – 0.09 % and precision of 0.035 – 0.09 % (Methods).

40 µm displacement applied to the sample resulted in a bulk IVD compression of 2.2 ± 0.7 %, calculated from the change in distance between the VEPs (Methods). In the caudal VEPs, 40 µm displacement of the sample generated a 1^st^ principal (tensile) strain of 0.6 ± 0.3 %, 3^rd^ principal (compressive) strain of 0.7 ± 0.3 %, and maximum shear strain of 0.7 ± 0.3 %. In the cranial VEPs, the resulting 1^st^ principal (tensile) strain was 0.7 ± 0.4 %, 3^rd^ principal (compressive) strain was 0.8 ± 0.4 %, and maximum shear strain was 0.8 ± 0.4 %.

The direction of 1^st^ (tensile) and 3^rd^ (compressive) principal strains with respect to the loading axis of the joint was measured as an angle between 0° (parallel to axis) and 90° (normal to axis). In the caudal VEPs, 1^st^ principal strains were at 59° ± 0.9° and 3^rd^ principal strains were at 54 ± 1.8° to the loading axis. In the cranial VEPs, 1^st^ principal strains were at 59° ± 0.9° and 3^rd^ principal strains were at 54 ± 2.6° to the loading axis.

The impact of radiation dose on measured strains was investigated (Supplementary Information, Figure S6). No statistically significant changes in mean VEP strain were seen with increasing radiation dose for cumulative dose levels in the IVD equivalent to 27, 53, and 80 kGy (Figure S6a, b). There was no increase in the number of local strain values exceeding 10 % strain with increasing radiation dose (Figure S6c, d). This indicates that radiation dose had no significant impact on the results of this study.

### Thickening of the peripheral endplate regions enables optimal load transfer from the IVD

To evaluate regional variations in VEP calcified tissue architecture that may modify load transfer from the IVD, sCT and DVC datasets for each endplate (*n* = 16) were subsampled into 10 standardised cubic sub-regions (side length 325 µm) for statistical analysis of structure and strain (Figure 3a). Each of the 8 whole IVD samples (2 endplates per sample), were treated as independent to account for variation in loading, spinal level, and image dynamics. Repeated measures correlation, which determines the common *‘within sample’* association for paired measures^58^, was used to test for relationships between radial location and microarchitecture, GSV, and local mechanical strain in the sub-regions (Fig. 3b). Holm-Bonferroni correction was used to deal with familywise errors^59^, and relationships were deemed significant at the 5 % level. This revealed a decrease in compressive (magnitude of 3^rd^ principal strain, R = −0.48, p < 0.001), tensile (1^st^ principal strain, R = −0.29, p < 0.001), and shear strain (maximum shear strain, R_rm_ = −0.41, p < 0.001) with radial distance from the endplate centre. On average, compressive strain in central endplate regions (Figure 3ai region 1) was 0.8 ± 0.5 % higher than the outer endplate regions (Figure 3ai regions 4,7,10), maximum shear strain was 0.6 ± 0.4 % higher, and tensile strain 0.4 ± 0.3 % higher after 80 µm cumulative displacement to the sample. This increase in local endplate load induced strain below the NP agrees with previous theories that the central endplate region experiences higher strains in vivo ^52,57^, and is clearly visible in lower resolution DVC strain maps (Figure S5).

**Figure 3.**
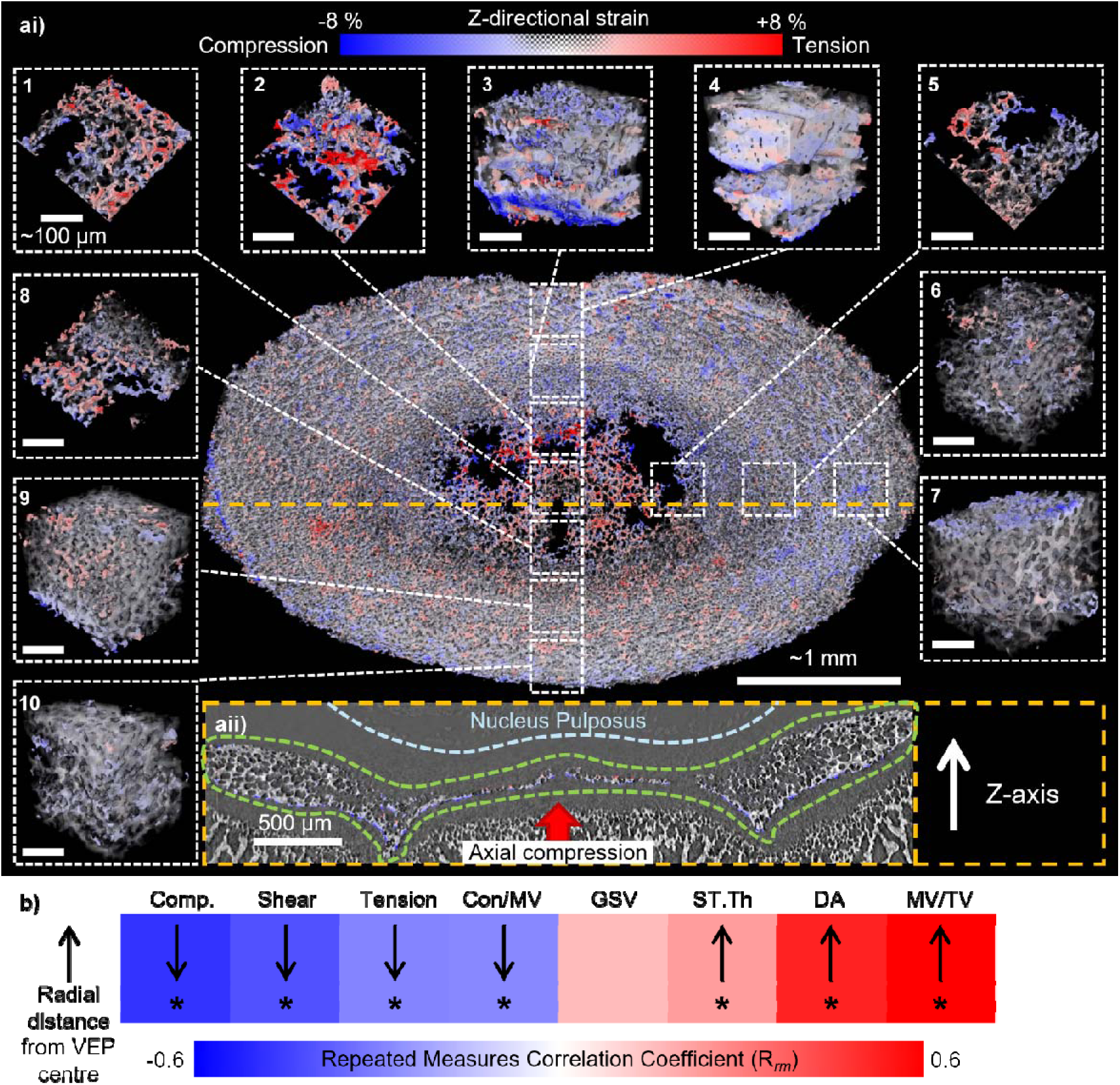
Thickening of the peripheral endplate regions enables optimal load transfer from the IVD. ai) 3D renderings of z-directional strain in a L2-3 cranial endplate. Regions used for statistical analysis are shown in dashed boxes, with scalebars representing ∼100 µm; 1 = centre, 2 = anterior inner, 3 = anterior middle, 4 = anterior outer, 5 = lateral inner, 6 = lateral middle, 7 = lateral outer, 8 = posterior inner, 9 = posterior middle, 10 = posterior outer. aii) Greyscale image with z-directional strain colourwash showing a slice through the orange dashed line on the 3D rendering in i, the dashed green line outlines the vertebral endplate. b) Heatmap of repeated measures correlation coefficient (Rrm) values for mean compressive strain (Comp.), mean maximum shear strain (Shear), connectivity to bone volume ratio (Conn/MV), mean tensile strain (Tension), mean greyscale value (GSV), mean septal/trabecular thickness (ST.Th), degree of anisotropy (DA), and mineralised tissue volume fraction (MV/TV), versus radial distance from the centre of the endplate (distance defined categorically as centre, inner, middle, or outer); downwards arrows represent a negative and upwards arrows a positive correlation; * represents statistical significance at the 5 % level after Holm-Bonferroni correction (see Table S5 and Figures S6 and S7 for full statistical analysis);

Mineralised tissue volume fraction (MV/TV) increased with radial distance from the endplate centre (R_rm_ = 0.76, p < 0.001), corresponding to an increase in thickness of the endplate towards the periphery. The degree of anisotropy was greater towards the outer endplate (R_rm_ = 0.52, p < 0.001), potentially due to the highly directional forces applied by the AF. Connectivity to mineralised tissue volume ratio (Con/MV) decreased (R_rm_ = −0.28, p < 0.001) and septal/trabecular thickness (ST.Th) increased (R_rm_ = 0.23, p < 0.01) towards the peripheral endplate, indicating a transition from calcified cartilage to bone, which can be distinguished by their resident cellular components visible in our sCT images. In bone, osteocytes reside in lacunae approximately 5 µm in diameter, in agreement with literature^60^, which reside in trabeculae 30 – 120 µm in thickness (Figure 2g). Calcified cartilage contains hypertrophic chondrocytes 20 – 50 µm in size embedded in mineralised lacunae^61^, creating a highly porous structure with mineralised cartilage septa 8 – 30 µm in thickness (Figure 2g). Accurate segmentation of bone and calcified cartilage in the VEPs would be extremely difficult and is not attempted in this study. Instead, microarchitectural values are used to indicate the predominant tissue type in a region, with higher values of ST.Th and MV/TV combined with lower values of Con/MV showing that a region predominantly contains bone.

### Spatial mapping of VEP structure to load-induced strain reveals tension and shear as potential drivers of the cartilage to bone transition

To interrogate relationships between microarchitecture, relative TMD (approximated by GSV), and local mechanical strain, repeated measures correlation^58^ was again used to test for links between the mean values for 160 endplate regions across 8 spine segments (Figure 4a). The generation and quantification of local strain induced by physiologically relevant whole joint loading enables an examination of how the structure of a region contributes to its ability to resist load, and also how the local strains in a region may impact the formation of mineralised tissues.

**Figure 4.**
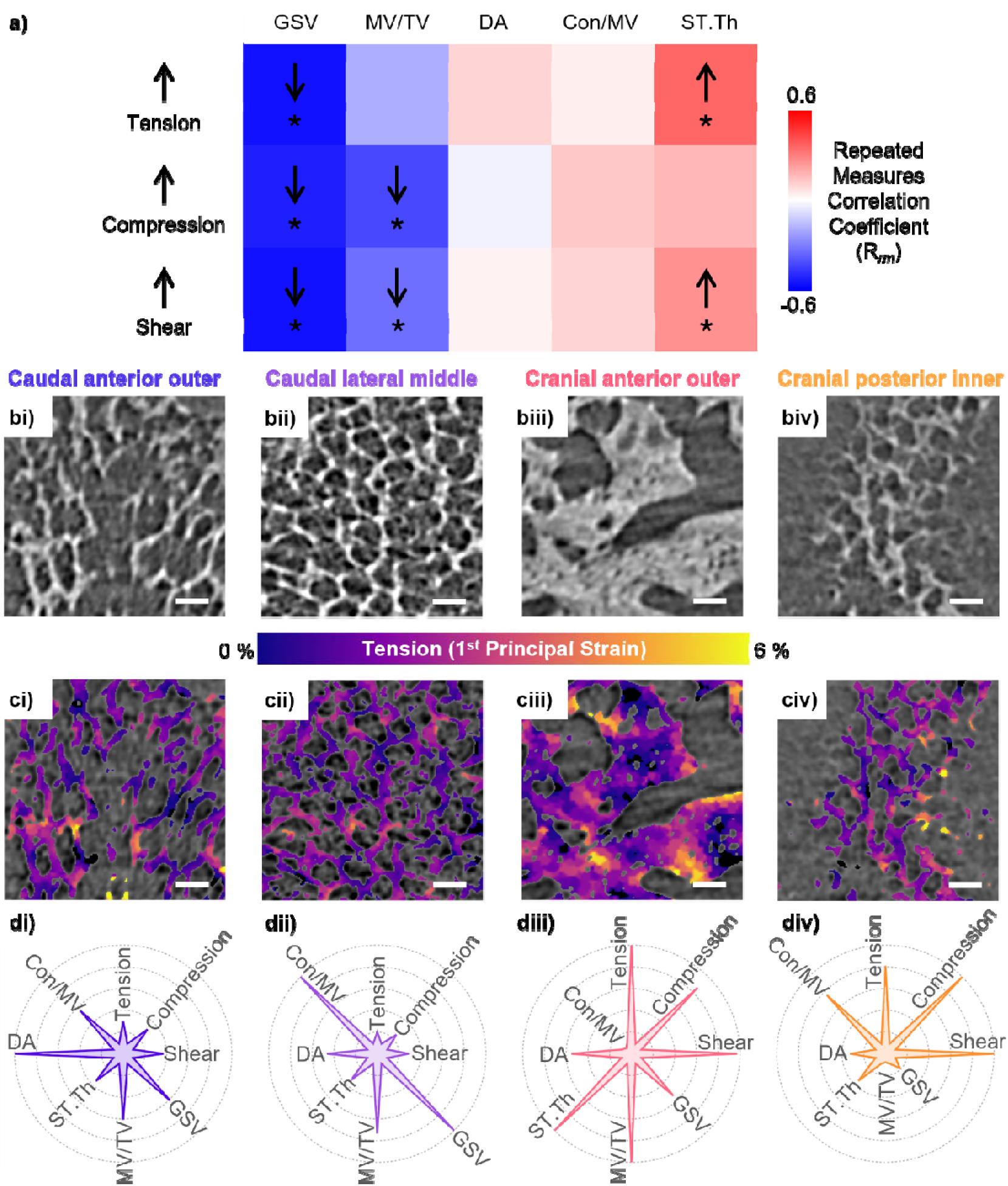
Spatial mapping of VEP structure to load-induced strain reveals tension and shear as potential drivers of the cartilage to bone transition. a) Heatmap of repeated measures correlation coefficient values for image greyscale value (GSV), mineralised tissue volume fraction (MV/TV), connectivity to mineralised tissue volume ratio (Con/MV), degree of anisotropy (DA), and mean septal/trabecular thickness (ST.Th) versus tensile strain (Tension), compressive strain (Compression), and maximum shear strain (Shear); * represents statistical significance at the 5 % level after Holm-Bonferroni correction, full statistical results given in Table S5 and Figures S8 and S9; b) Greyscale image slices, c) tensile strain fields (compressive strain fields shown in Figure S11), and d) radar plots of relative strains and structural parameters between 4 regions of a lumbar 4-5 sample: i) the caudal anterior outer, ii) the caudal lateral middle, iii) the cranial anterior outer, and iv) the cranial posterior inner region. Scale bars represent 50 µm. Values used to generate radar plots available in Table S6.

Negative correlations were seen between GSV and local mechanical tensile (R_rm_ = −0.56, p < 0.001), compressive (R_rm_ = −0.53, p < 0.001), and shear strain (R_rm_ = −0.56, p < 0.001), agreeing with previous studies showing increased stiffness with higher mineralisation^62^. MV/TV negatively correlated with local mechanical compressive (R_rm_ = −0.43, p < 0.001) and shear strains (R_rm_ = −0.34, p < 0.001), and ST.Th showed positive correlations with local tensile (R_rm_ = 0.35, p < 0.001) and shear strain (R_rm_ = 0.25, p < 0.01).

These correlations reveal two predominant structural types in high strain regions. First, a low TMD calcified cartilage region (identified by the presence of chondrocyte lacunae, Fig. 4biv) with low MV/TV and thin septa found in the central and inner endplate regions adjacent to the NP, which we speculate experiences elastic deformation under loading, leading to high local strains. The second type of high strain region contains trabeculae, high MV/TV, and low Con/MV, indicating that these regions have undergone endochondral ossification to form bone (confirmed by the presence of osteocyte lacunae, Figure 4biii).

### Correlative imaging and diffraction enables colocalisation of endplate microarchitecture with nanoscale mineral crystal properties

To enable colocalization of microarchitecture in VEP calcified cartilage and bone compartments with these nanoscale mineral crystal properties, consecutive, spatially correlated sCT and WAXD raster scans (Fig. 5a-c) of partial IVDs was performed at the DIAD beamline, DLS ^54^ (Methods). Physiologically relevant conditions were applied to fresh-frozen quarter IVD segments, with samples held under a tensile prestrain to simulate AF native tensile collagen prestrain and hydration maintained. The spatial resolution of sCT images was 3.2 µm, calculated from Fourier shell correlation analysis (Figure S14). Axial and transverse d-periods were derived from the positions of the (002) and (310) WAXD peaks respectively (Figure 5d,e), and mineral crystallite length and width were calculated from the corresponding peak widths^13^ (Figure 5f-g, Methods). Crystallite dimensions were 11.4 ± 1.7 nm (length) and 1.23 ± 0.13 nm (width), consistent with prior results in young, mineralised tissues^11,63^. The (002) d-period varied between 3.42 and 3.44 Å, corresponding to a variation in mineral prestrain along the c-axis of 0.6 %. The (310) d-period varied between 2.26 and 2.28 Å, corresponding to a variation in mineral prestrain along the a-b-axis of 0.9%. Assuming a mineral Youngs modulus of ∼100 GPa, these prestrain variations result in 600 - 900 MPa prestress in the mineral.

**Figure 5.**
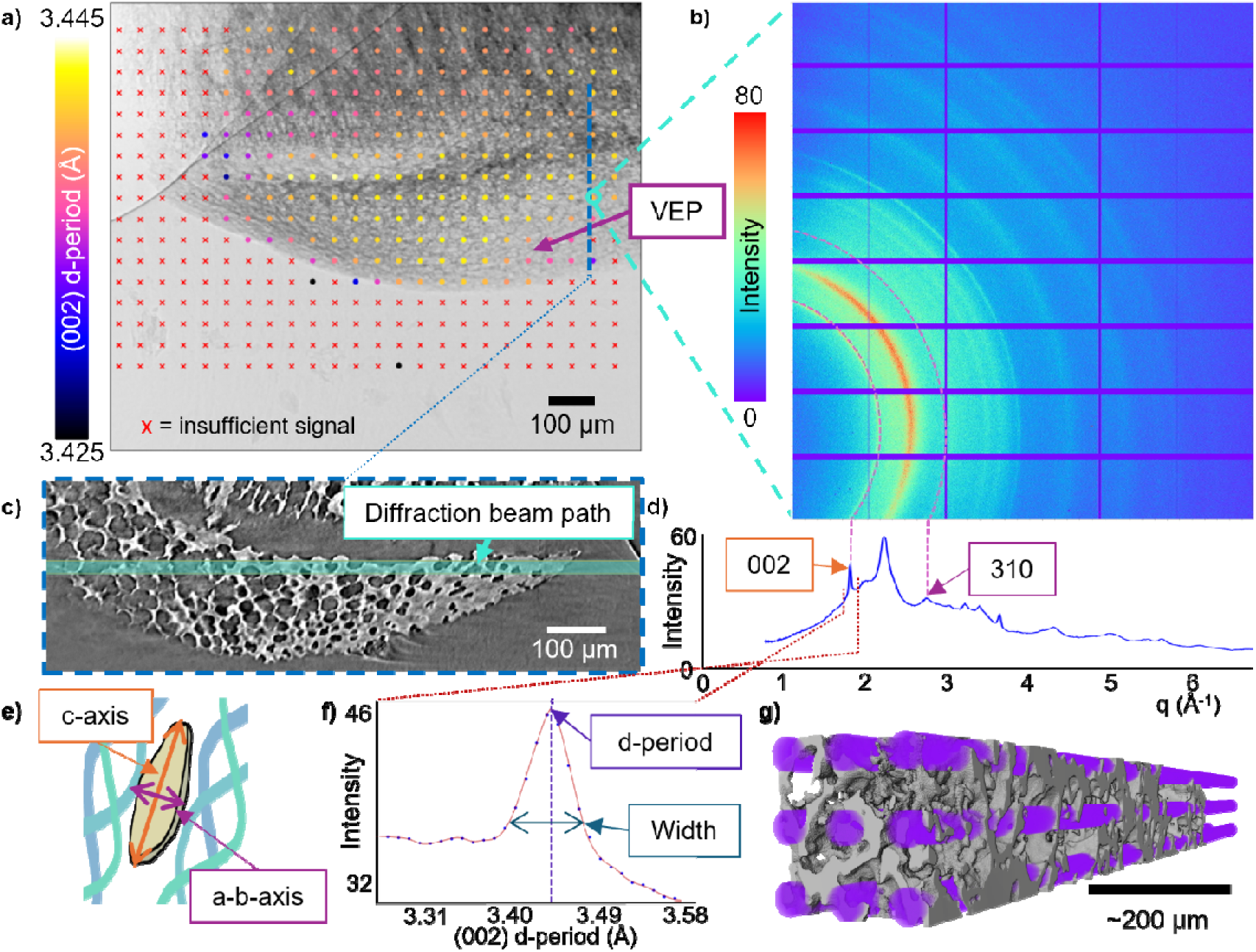
Correlative imaging and diffraction enables colocalisation of endplate microarchitecture with nanoscale mineral crystal properties. a) Map of hydroxyapatite (002) lattice spacing on an X-ray projection of a quarter IVD sample held under tension. b) WAXD intensity pattern from the location highlighted in a), only ∼1/4 of the full diffraction rings are visible due to the need to place the WAXD detector off-centre to allow for placement of the imaging camera for sCT (Methods); c) sCT image of the endplate showing the location of the diffraction beam path that produced the WAXD pattern in b), the location of the reconstructed slice is shown as the dashed blue line on the projection in a); d) WAXD pattern in b) integrated over q; e) illustration of a HAP platelet surrounded by collagen, with c and a-b axes labelled; f) HAP (002) peak plotted in d-space, with d-period and width labelled; g) 3D rendering of calcified tissue structure (grey) and diffraction beam paths (purple) for one region used for microstructural evaluation and statistical analysis.

### Bone is under greater compressive prestrain and contains narrower crystallites compared to calcified cartilage

New insights into relationships between microscale VEP mineralised tissue architecture and nanoscale crystallite structure/ prestrain were therefore examined by analysing mean values of regions of interest comprising nine WAXD beam paths (Fig. 5g); a total of 194 regions were taken across 8 quarter IVD samples from 4 rat lumbar spines for statistical analysis (Figures S15 and S16). Each quarter IVD sample was treated as independent and repeated measures correlation used to test for common - within sample - relationships (Figure 6a). This showed that both (002) and (310) prestrain (d-period) in the mineral was higher (more tensile) in regions with greater mineralisation density/ GSV (R_rm_ = 0.35, p < 0.001; R_rm_ = 0.24, p < 0.01).

**Figure 6.**
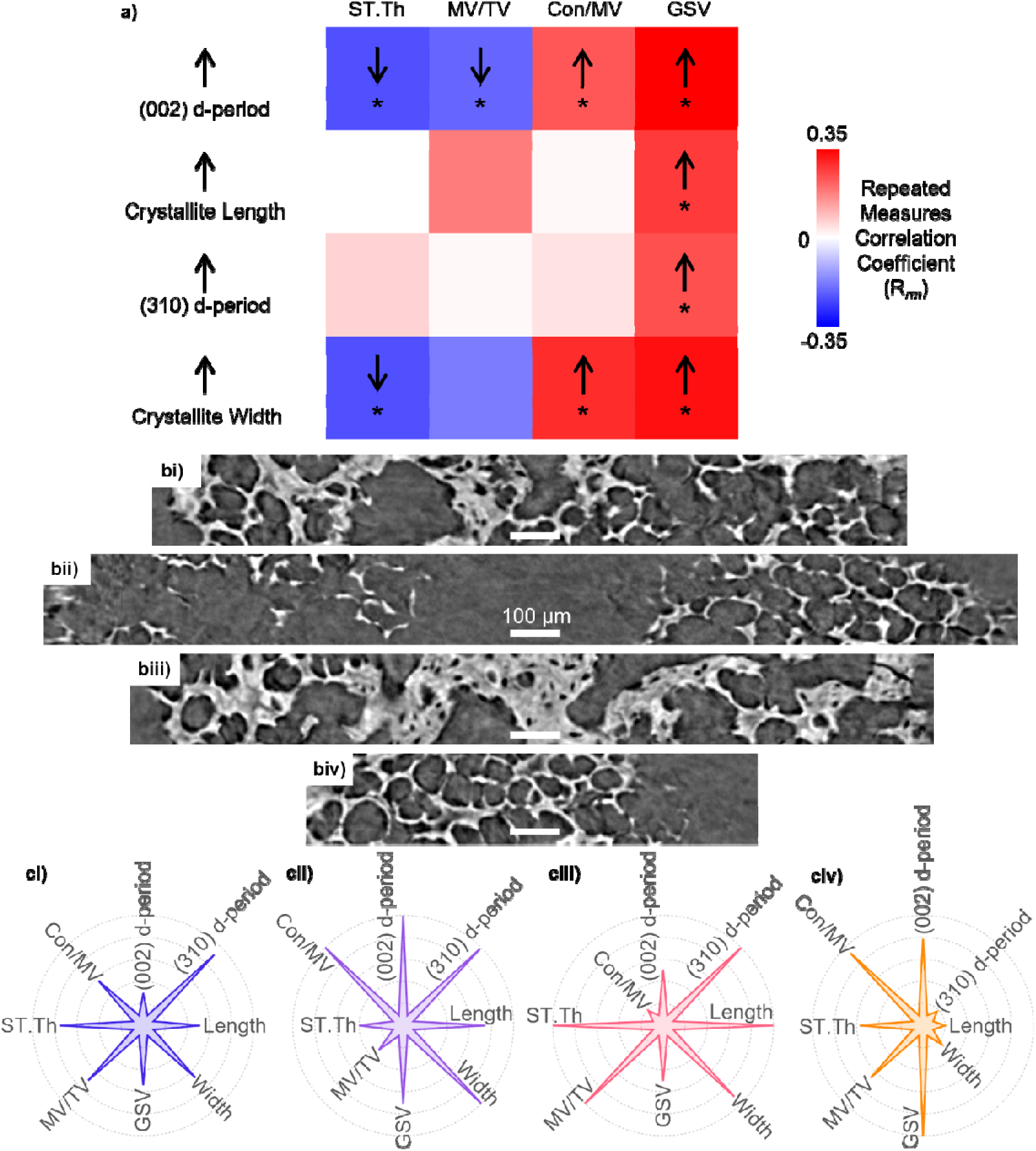
Bone is under greater compressive prestrain and contains narrower crystallites compared to calcified cartilage. a) Repeated measures correlation coefficient matrix for mean septal/trabecular thickness (ST.Th), mineralised tissue volume fraction (MV/TV), connectivity to mineralised tissue volume ratio (Con/MV), and image greyscale values (GSV) versus mineral (002) lattice spacing ((002) d-period), mineral crystallite length (Length), mineral (310) lattice spacing (310 d-period), and mineral crystallite width (Width); * represents statistical significance at the 5 % level after Holm-Bonferroni correction, full statistical results are presented in Table S8 and Figures S15-S17. b) Greyscale cross-sections from four VEP regions used for statistical analysis from one sample showing microscale structural variation in the VEP, i) contains a mixture of calcified cartilage and bone, ii) contains sparse calcified cartilage, iii) consists of mostly bone, and iv) contains calcified cartilage, scalebars represent 100 µm. c) Radar charts displaying the relative values of WAXD and sCT based measurements from the four regions shown in b (the values used to generate the radar charts is presented in Table S9).

(002) d-period was lower in regions where structural organisation indicates the presence of bone, decreasing with MV/TV (R_rm_ = −0.21, p < 0.01) and ST.Th (R_rm_ = −0.24, p < 0.001) and increasing with Con/MV (R_rm_ = 0.23, p < 0.01). This indicates a greater compressive prestrain in regions that have undergone endochondral ossification (such as the regions shown in Fig. 5bi and iii), and that calcified cartilage is under tensile prestrain relative to bone. This supports the intriguing possibility that the local mechanical equilibrium of the mineral crystallites is functionally distinct between calcified cartilage and bone, possibly due to differences in organic composition.

Positive correlations were seen between GSV and mineral crystallite length (R_rm_ = 0.33, p < 0.001) and width (R_rm_ = 0.27, p < 0.001), confirming that more mature mineral has larger crystallites. Crystallite width decreased with ST.Th (R_rm_ = −0.24, p < 0.001) and increased with Con/MV (R_rm_ = 0.29, p < 0.001). This indicates that regions containing bone have narrower crystallites than regions containing calcified cartilage. Positive correlations were seen between (002) d-period and crystallite length (R_rm_ = 0.34, p < 0.001), and (002) d-period and crystallite width (R_rm_ = 0.44, p < 0.001), implying that longer, wider crystallites are formed when placed under tensile strain along the fibrillar axis, potentially due to greater alignment of gap-overlap channels in the fibril under greater tensile strain.

## DISCUSSION

This study represents a step-change advance in our understanding of the multiscale origins of IVD biomechanical function, with implications for the investigation of ageing and degenerative joint diseases such as IVD degeneration and osteoarthritis in future studies. The combined analysis of load-induced tissue strains using DVC and molecular-scale mineral prestrain using WAXD, has revealed the important role of mineral prestrain in forming the complex biomechanical strain patterns underpinning whole spinal joint level mechanics (Figure 7). At the whole joint scale, the decrease in overall VEP thickness below the NP enables central endplate bulging and accumulation of high tissue level strains in these regions (Figure 7). At the microscale, higher mineralised tissue volume fraction and trabecular thickness below the AF helps resist tensile and shear strains, with these regions showing a higher degree of anisotropy (Figure 7); we hypothesise that the preferential orientation of mineralised structures in these regions helps resist highly directional tensile loads from collagen fibres in the AF. The direction of tensile strains in the VEPs was 59° to the loading axis, and compressive strains 54°, demonstrating how the IVD converts axially applied loads to more radial and circumferential forces in the endplates, which we hypothesise helps prevent vertebral fracture in axial loading environments. These micro- to whole-joint-scale structure-strain relationships were conserved across all spinal levels investigated (thoracic 10 to lumbar 5). At the nanoscale, differences in prestrain and crystallite size in bone and calcified cartilage show interesting links to the strain patterns observed in whole VEPs (Figure 7). It should be noted that due to the nature of the investigation performed, only association, not causation, can be drawn from this study. Future studies should aim to test the hypotheses discussed below.

**Figure 7.**
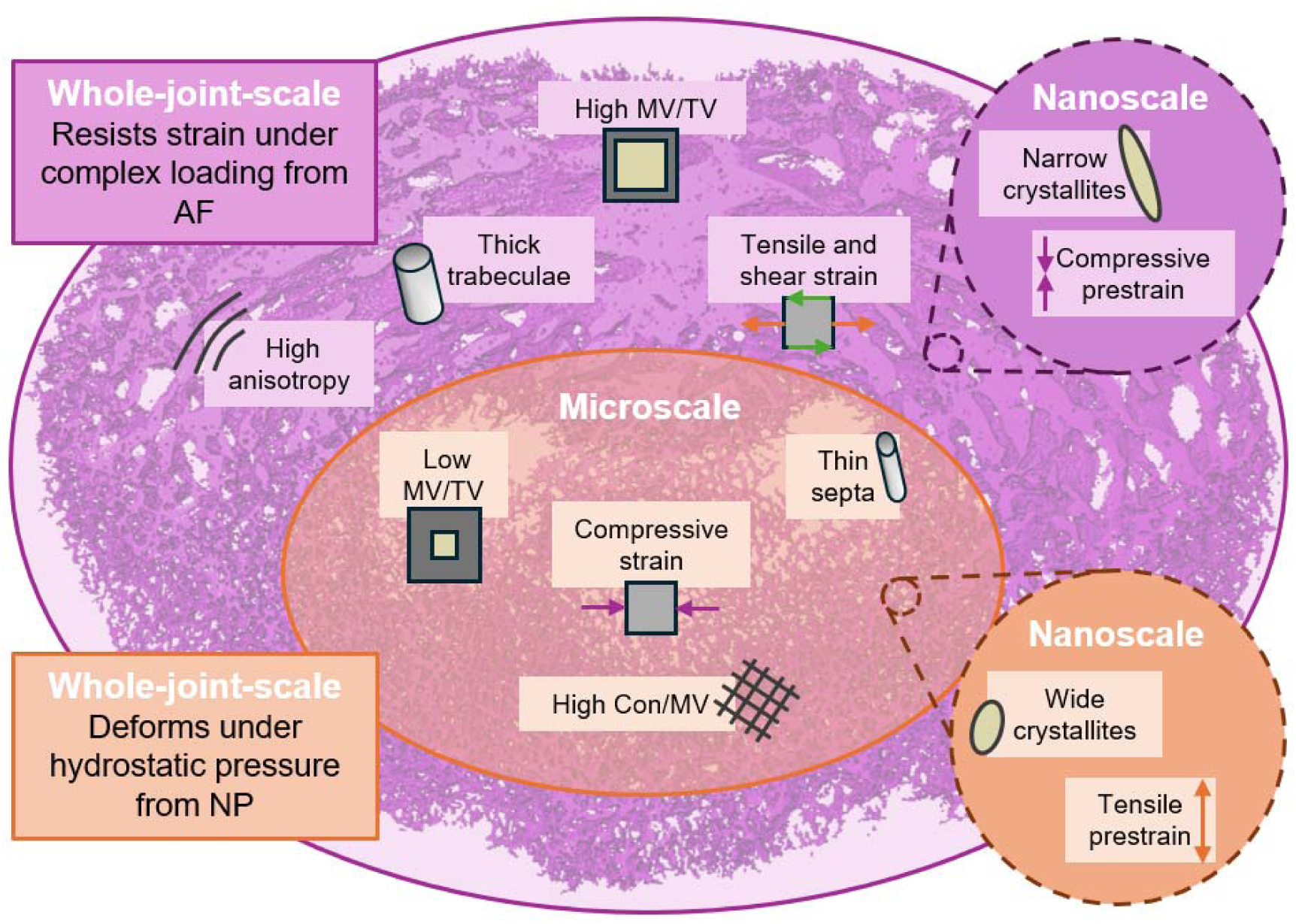
Summary of the differences in loading environment, structure, and prestrain between the central and peripheral vertebral endplates. At the whole joint scale, the peripheral endplate resists complex loading from the annulus fibrosus (AF); at the microscale the peripheral endplate has high anisotropy, thick trabeculae, high MV/TV (mineralised tissue volume fraction), and experiences high tensile and shear strains; at the nanoscale the peripheral endplate contains narrow mineral crystallites which are under relative compressive prestrain. The central endplate deforms under hydrostatic compressive loading from the nucleus pulposus (NP); at the microscale the central endplate has low MV/TV, thin septa, high Con/MV (connectivity to mineralised tissue volume), and experiences high compressive strains; at the nanoscale the central endplate contains wide mineral crystallites which are under relative tensile prestrain.

DVC found higher load induced tensile and shear strains in regions containing bone (Figure 4biii, 7), revealing that local mechanical strain may play a driving role in the cartilage to bone transition, providing support for the theory that hydrostatic compression inhibits endochondral ossification^19^. The microscale resolution and nanoscale precision of DVC achieved in this study enables analysis of the local cellular mechanical environment under whole-joint loading, demonstrating the potential of DVC as a tool to bridge the gap between cell mechanobiology and whole joint biomechanics.

WAXD analysis provided nanoscale information on the structure and prestrain state of the mineral component of the extracellular matrix. Regions containing bone (Figure 6biii, 7) showed greater mineral *compressive* prestrain which, we hypothesise, both increases VEP toughness in these regions due to mineral compaction, enabling them to resist compression with minimal deformation, and allows for greater levels of tensile strain to be engendered before failure.

In contrast, VEP regions adjacent to the NP (Fig. 4biv) were shown to experience the highest tissue-level compressive strain, agreeing with theories that the endplates bulge under hydrostatic pressure from the NP^52,57^. Our WAXD analysis and quantification of mineralised tissue architectural features in these regions suggest that the mineral is instead under *tensile* prestrain relative to the bone regions mentioned previously, which may provide VEP calcified cartilage with a greater level of elastic compressibility under loading, enabling central endplate bulging to help prevent NP over-pressurisation which would otherwise lead to herniation^52^. This observation of prestrain in the central VEPs is supported by experimental results in human IVDs, where the removal of the nucleus pulposus resulted in endplate bulging^64^. We propose that regional variations in mineralised tissue prestrain may play a vital role in the optimal mechanical function of musculoskeletal joints, and the role of prestrain in conditions such as osteoarthritis, osteoporosis, and IVD degeneration should be further investigated.

This variation in mineral prestrain in calcified cartilage and bone has an interesting correlation with similar findings of higher collagen fibril prestrains (at one step higher in the hierarchical structuring) in calcified cartilage than subchondral bone, measured from small-angle X-ray scattering (SAXS) across bovine metacarpophalangeal joints^29^. These fibrillar differences were interpreted as an adaptation in cartilage allowing for elastic deformation under compression – deforming from an initial relative tensile state to an effective zero load state when compressed. Here, the origin of the lowered mineral prestrain in bone could be explained if the collagen fibril-mineral interface is tighter than in calcified cartilage, leading to greater compressive stress exerted on the mineral platelets by the surrounding collagen (possibly due to a more organised mineral-collagen interrelationship). It has been shown that intrafibrillar mineralisation generates collagen fibrillar contraction, but extrafibrillar mineralisation does not^33^; in the context of our results, this would imply that bone has more intrafibrillar mineralisation whereas cartilage mineralisation is more extrafibrillar. In this regard, these prestrain differences could also be related to the differing proteoglycan content of bone and cartilage^8,65,66^. Aggrecan, the large aggregating proteoglycan, in cartilage is known to generate a swelling pressure within the tissue, placing collagen fibres under pre-tension, and some remnants of this tensile prestrain may remain in the tissue after mineralisation, resulting in a tensile mineral prestrain in calcified cartilage relative to bone. Prestrain differences could also be the result of the different cell populations in these two tissues, with different cell types actively generating differing amounts of tension in the extracellular matrix. Differing mineral substitutions between calcified cartilage and bone^8^ could also result in prestrain differences by disrupting the lattice. An increase in d-period was seen with mineralization density, indicating a reduction in compressive prestrain in the mineral. This aligns with the concept of load sharing between mineral and collagen, and axial fibrillar contraction ^16^, which we propose leads to greater compressive load on the mineral in regions of lower TMD. Here, we have demonstrated using sCT and DVC how these variations in prestrain can be related to whole joint mechanics, furthering the hypothesis that - at the molecular level - relative tensile prestrain in the stiff constituents of the calcified cartilage extracellular matrix enables elastic deformation under macroscale compressive load, whilst compressive prestrain in bone confers resistance to complex loading environments involving tension and shear.

We also observe that regions containing bone have narrower mineral crystallites than calcified cartilage regions (Figure 7), suggesting differences in mineralisation between these two tissue types. Calcified cartilage contains more extrafibrillar space than bone^63^ and contains mostly type II collagen rather than majority type I^29^ which may exert a role in the formation of bone apatite^12^. We speculate that the crystallites in mineralised cartilage may be wider due to their extrafibrillar, as opposed to intrafibrillar, bone-like mineralisation^11,67^. Mechanically, computational modelling has shown that plastic spreading of crystallites first occurs in the extrafibrillar space, followed by the intermolecular space^68^, so differences in mineral location between calcified cartilage and bone would result in differing material properties. Osteoblasts in bone and chondrocytes in cartilage have different directional preferences for the initiation of mineralisation by matrix vesicle secretion^69^, and we hypothesise that this may also play a role in the differences in crystallite shape observed in this study. Studies have shown that crystallites with larger aspect ratios increase the stiffness, strength, and toughness of mineralised tissues^70,71^, and the wider mineral crystallites in calcified cartilage would have a lower aspect ratio. The variation in prestrain and crystallite shape seen here may explain why bone is stiffer and harder than calcified cartilage even when their mineral content is similar^7^, and supports the notion that calcified cartilage acts as an important transition layer in diarthrodial joints, reducing the strains generated at hard-soft tissue boundaries^6,29,63^. Changes in mineralisation and mineralised tissue structure are associated with multiple pathologies, including osteoarthritis ^8,15^ and IVD calcification^72^, and the combination of methods presented in this study should be used to investigate these conditions in future studies.

This study demonstrates that regional variations in multiscale structure and strain at the joint articulating surface contribute to overall mechanical performance. These regional variations should be considered when designing orthopaedic implants and biomaterials, for example gradient scaffolds. Gradient scaffolds for osteochondral repair improve repair and regeneration compared to homogeneous scaffolds but still require improvement for clinical translation^73^. Improvements in manufacturing methods such as 3D printing means that producing biomaterials with varying microarchitectures is becoming more common^74^, and the microarchitectural relationships uncovered here should be used to inform the design of novel biomaterials. Nanostructured materials are showing huge promise for orthopaedic biomaterials development^75^, we propose that the introduction of a nano-crystallite size/ shape gradients and molecular level prestrain may improve the functionality of biomaterials for orthopaedic applications.

The rat model used herein is considered representative of young, healthy IVD, and was chosen primarily based on size with respect to the imaging field of view available in this study, and because of the common use in preclinical research. Rat lumbar disc geometry resembles human discs, so is suitable for the investigation of axial mechanics^76^. The young age (8-weeks) of the animals was chosen to enable investigation of mineralised structures that form during the growth stage, and it should be noted that the structure-strain relationships uncovered in this study may change with age as mineralised structures further adapt to their mechanical environment. Future studies should investigate how these relationships change with age and degenerative conditions including osteoarthritis and IVD degeneration.

Limitations of this study include scan times, radiation dose, and the need to perform two separate experiments. The 3-minute scan times used for whole IVD *in situ* imaging meant that truly dynamic imaging was not possible. The quasi-dynamic regime used was able to capture load induced strains after full stress relaxation, meaning a significant proportion of strain was taken up by the soft tissues of the IVD. Dynamic mechanical responses occurring during and immediately after loading could not be captured in this study, and it is likely the mineralised tissues experience higher strains immediately after the load is applied, making our results a lower bound. Scan times on the order of seconds have been achieved to dynamically investigate cartilage biomechanics^41^, and future studies of the IVD should look at utilising similar fast imaging set-ups. The accuracy of the load cell used in this study (±1 N) was insufficient for the collection of reliable force data and means there was likely variability in sample preload, this was accounted for statistically by treating each mechanical experiment, consisting of preload plus three displacement steps, as independent.

Radiation dose can cause a change in the elastic properties of bone and induce microcracking^77^, so attempts were made to reduce radiation dose during all experimental procedures. The applied dose per image during *in situ* sCT was 27 kGy, below the 35 kGy often quoted as safe for bone samples^78^, and studies on cartilage biomechanics have shown no clear radiation dose effects on *in situ* mechanics at doses up to 300 kGy^41^. Here, no changes in measured strain values in the VEPs were seen with increasing radiation dose (Figure S6), so it can be concluded that radiation dose did not have a statistically significant impact on the results of this study.

The WAXD patterns collected in this study covered only a quarter of the azimuthal range, preventing the analysis of crystallite orientation. This study used 2D WAXD analysis, but 3D diffraction techniques can provide a fuller picture of how nanoscale structure and prestrain change throughout complex sample geometries^46^. The tensile preload applied to partial IVD samples during correlative sCT and WAXD was designed to mimic physiological conditions, but this may have resulted in some load-induced strain in the mineral as opposed to purely prestrain. However, the maximum load induced strain seen in the mineral portion of bone during elastic loading is around 0.2 %^17,78^, so the ∼1 % variation in mineral strain seen in this study must be predominately due to prestrain effects. This study used two separate beamlines to achieve both *in situ* sCT of whole joint loading and correlative sCT and WAXD; ideally, a beamline set-up enabling simultaneous acquisition of both datasets should be used.

## CONCLUSIONS

Combined sCT, DVC, and WAXD analyses have enabled whole-joint to nanoscale structural quantification of murine VEPs, showing significant radial variation, transitioning from a thin layer of calcified cartilage centrally, adjacent to the NP, to thicker peripheral bone structures close to the AF (Fig. 7). DVC enabled quantification of 3D strain fields in VEP microarchitectures, revealing the presence of higher load-induced strains in the central VEPs adjacent to the NP. Regions containing bone had higher loading-related tissue level tensile and shear strains, potentially demonstrating the instructive role of mechanical loading in the cartilage to bone transition, with complex loading environments involving tension and shear leading to bone formation, and hydrostatic compressive environments remaining as cartilage. Correlative sCT and WAXD exposed divergent mineral prestrain in bone and calcified cartilage, with calcified cartilage under tensile prestrain relative to the compressive prestrain of bone regions (Fig. 7). Regions containing bone had narrower crystallites compared to calcified cartilage, which likely indicates that cartilage undergoes more extrafibrillar mineralisation. Mineral crystallites were both longer and wider when under relative tensile prestrain. Linking these nanoscale results back to the structure and function of the VEPs, we hypothesise that compressive prestrain and a higher crystallite aspect ratio in the bone around the peripheral VEPs aids in resisting complex loading from the AF, whereas tensile prestrain and a low crystallite aspect ratio in the calcified cartilage below the NP enables its elastic deformation under hydrostatic compression and prevents NP over-pressurisation.

Our data demonstrate the potential of *in situ* multimodal X-ray probing for multiscale biomechanics and mechanobiology. Each modality offers a unique complementary dataset, allowing a holistic approach to interpreting biomechanics from the whole joint to molecular scales. Future applications of these techniques have the potential to unravel the biomechanical underpinnings of biomineralization, growth, aging, and the effects of pathologies known to affect mineralised tissue structure, including osteoarthritis and osteoporosis. The multiscale structure-function relationships uncovered here should be used to inform the design of novel biomaterials.

## METHODS

### Animals and sample preparation

Male 8-week-old Sprague Dawley rats kept in controlled conditions (12:12 hour light:dark cycles, 22 °C and *ad libitum* access to food and water) at the University of Manchester Biomedical Sciences Facility with all tissue collection procedures performed in accordance with the UK Animals (Scientific Procedures) Act 1986, local regulations set by the UK Home Office with institutional approval obtained from the University of Manchester Animal Welfare and Ethics Review Board under the Home Office Licence (#70/8858 or I045CA465). Whole spines were dissected and snap frozen in liquid nitrogen prior to storage at −80°C. Within 24 hours of *in situ* mechanical testing, samples were thawed at room temperature before further dissection.

For *in situ* sCT of whole IVD, 9 individual spinal units across 3 spines (from vertebral levels thoracic 9 to lumbar 6) consisting of an IVD with half vertebrae either side and posterior elements removed, were used for analysis. Removal of posterior elements was required to enable visualisation and isolation of the IVD. Dissections were performed using a high precision diamond cutting blade (Accumtom-50; Struers, UK). Sample damage was minimised during dissection through the use of water cooling, a slow feed speed (0.05 mm/s) and 1800 rpm. Spinal level was recorded in terms of vertebral number either side of the IVD for each sample (Table S1).

For correlative imaging and diffraction experiments, spinal units from spinal levels lumbar 3 to lumbar 6 were further dissected into quarters along the spinal axis, resulting in 8 partial vertebra-annulus-vertebra samples from 4 spines used for analysis (Table S7). This was done to ensure samples fit within the smaller imaging field of view at the DIAD beamline (1.38 x 1.16 mm).

### In situ mechanical testing

Samples were mounted in a Deben CT500 (Deben, UK) rig containing a 100 N loadcell (accuracy ±1 % of full scale) and custom-built open frame design consisting of an aluminium alloy base cover and two carbon fibre pillars bridged by an aluminium alloy crossbar. The rig had a 10mm linear extensometer for position readout, with a 3 μm resolution, and linearity 1% of full scale. The open frame design gave easy access to the samples during alignment and rehydration and reduced the amount of material in the path of the diffraction beam for dual imaging and diffraction experiments. Custom 3D printed (Acrylonitrile Butadiene Styrene, ABS) sample holders were used for sample mounting.

For in situ sCT of whole IVDs, spinal units were fixed to the bottom 3D printed compression platen using superglue (Vetbond, 3M) and left free at the top. Samples were placed inside polyimide tubing (diameter 6mm, wall thickness 75 µm) filled with phosphate buffered saline (PBS) to maintain sample hydration during scanning (Figure S1). Samples were held under cumulative axial compression, applied in the z-direction from beneath the bottom sample holder during imaging. A 1 N compressive preload was first applied, corresponding to approximately 40 % the animals body weight (260 ± 33 g), designed to prevent over swelling of the nucleus pulposus and generate a physiologically relevant level of intradiscal pressure by simulating super-incumbent body weight^57^. The first sCT image was taken 20 minutes after application of the preload to allow for stress relaxation and reduce motion artefacts acquired during scanning. Three further images were taken after applying successive 40 μm displacement steps. Peak and relaxed load readouts for each sample are given in Table S2. All loads and displacements were applied at a slow rate of 0.1 mm/minute to reduce the amount of stress relaxation after loading. The full force, displacement, time readout for one experiment (sample 5) is available in the supplementary dataset^79^.

For correlative imaging and diffraction experiments, partial vertebra-annulus-vertebra samples were fixed at both ends to 3D printed sample holders using superglue (Vetbond, 3M). Samples were placed inside polyimide tubing (diameter 4mm, wall thickness 76 µm) filled with PBS to maintain sample hydration during scanning (Figure S12). A 2.5 N tensile preload was applied before scanning to place annulus fibres into tension and prevent sample motion during scanning. sCT scanning was performed after 30 minutes stress relaxation, followed by WAXD scans.

### Phase contrast enhanced sCT imaging of whole IVD

sCT of whole IVDs was performed at the Diamond-Manchester Imaging Branchline I13-2 ^80^ at Diamond Light Source, UK using a pink beam with mean energy 27.6 keV. Low energy X-rays were removed using filters (Pyr. Graphite 0.28 mm + 1.06 mm, Al 3.2 mm, Fe 0.14 mm) to reduce radiation dose to the samples. Imaging setup (Sup. Fig. 9) was based on previous experiments ^56^, with the main changes being a more efficient scintillator, 500 μm LuAG SO14, and larger propagation distance of 0.5 m, allowing shorter exposure times to be used. 1800 projections with 0.1 s exposure time were taken per image volume using fly scanning, resulting in a scan time of 3 minutes and an applied X-ray dose of 27 kG, an 85 % reduction in dose compared to previous experiments ^55^, see supplementary information document for calculation. Images had a 1.6 µm voxel size and 4.2 mm x 3.5 mm field of view. Limited angle tomography artefacts resulting from the open frame mechanical testing rig were avoided through the use of carbon fibre pillars which were designed to be X-ray transparent, allowing collection of usable tomographic projections at all angles. One pillar covering the field of view results in a 22 % loss of intensity at the detector, and the pillars appear in 19 % of the collected projections (Figure S2). No correction for this loss of intensity was made during image reconstruction.

Tomographic datasets were reconstructed using the open-source pipeline Savu^81^. Firstly, projections were pre-processed using flat and dark field correction and lens distortion correction ^82^, followed by ring artefact removal ^83^. Datasets were then reconstructed using the filtered back projection algorithm ^84^ with Paganin phase retrieval ^85^ (δ/β value of 100) formatted in 16-bit tif stacks. An unsharp mask (radius = 2.5 pixels and weight = 0.9) was applied to the reconstructed datasets using ImageJ^86^ to reduce blurring associated with applying a Paganin filter. Example sCT images are available in the supplementary dataset^79^. The spatial resolution of the images was determined using Fourier shell correlation analysis (Figure S3).

### Point cloud generation for DVC

The DVC point cloud specifies the location of the centre of each subvolume used for tracking. Point clouds for the vertebral endplates were created using interactive thresholding and meshing in Avizo 2022.2 (Thermo Fisher Scientific, Waltham, Massachusetts, U.S.) (Figure 2h). Here, the vertebral endplates are defined as the epiphysis between the vertebra and the IVD (Figure 3aii). The interactive thresholding tool was used to create a binary image of the calcified tissue regions. The threshold value was chosen for each image individually, based on the value best able to separate mineralised tissue from the marrow space. The volume edit tool was then used to remove the underlying vertebral bone and ensure points could not be placed too close to the edge of the image or in regions severely affected by imaging artefacts. The Remove Small Spots module was used with size 2×10^7^ 3D to remove any disconnected regions from the binary image. The Remove Small Holes module was used with size 14000 3D to remove holes smaller than the DVC subvolume size. A Binary Smoothing was applied with a kernel size of 3 in all directions and threshold of 0.55, this slightly eroded the binary image away from the edges of the mineralised tissue and smoothed the image to improve meshing. A surface was then generated from the smoothed binary image and the surface simplified by increasing the minimum distance between nodes to 4.5 voxels (7.3 µm). A tetrahedral grid was generated from the surface and the nodes of the mesh used to specify locations of the points used for DVC (Figure 2h). This resulted in highly dense point clouds containing ∼1 million points per-endplate, resulting in a subvolume overlap of up to 80 %. VEP meshes and point clouds for sample 5 are available in the supplementary dataset^79^.

### Digital Volume Correlation

DVC was performed using the local approach with flexible point cloud specification^37,38^. Source code was supported by CCPi (Collaborative Computational Project in Tomographic Imaging, https://tomographicimaging.github.io/iDVC/, version 21.1.0). All DVC computations were run using spherical subvolumes with 12 degrees of freedom (translation, rotation, and 1^st^ order strain), using a zero-normalised sum-of-square difference (ZNSSD) function for optimisation. A subvolume diameter of 30 voxels (49 µm) was used with 2000 sampling points per volume. Greyscale value was interpolated within subvolumes at these sampling points using tricubic interpolation, enabling subvoxel displacement measurements. Lagrangian strains were calculated from the displacement field using polynomial fitting and differentiation^38^. A strain window of 25 points was used, resulting in a mean strain window radius of 21.4 μm, calculated as:

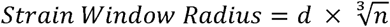

Where *d* is the mean distance between points in the point cloud (here 4.5 voxels) and n is the number of points used for polynomial fitting in the strain calculation (25). This resulted in a virtual strain gauge length of 91.5 μm, representing the span of points used in the strain calculation, calculated as the strain window diameter plus the subvolume diameter.

Accuracy and precision of the displacement and strain fields were calculated using a zero-strain analysis. Standard zero-strain analyses involve using two images taken sequentially of a sample without applying any load. This method can be problematic when imaging highly viscoelastic samples such as the intervertebral disc, as they are likely to have moved between scans, wrongly giving the impression of poor accuracy and precision. An alternative zero-strain method was used for this experiment, where one scan was taken with 3600 projections, which was then split into two sets of 1800 projections to be used for zero strain analysis^55^. This produces two independent scans taken at the same time, removing the potential impacts of viscoelasticity and providing a more robust method of determining true DVC accuracy. Images for zero-strain analysis were reconstructed without projections containing carbon pillars, using ∼1600 projections. This was done to prevent differences between the two images caused by the carbon pillars appearing in slightly different locations between the two scans. A point cloud was created for the cranial endplate of one lumbar 1-2 sample containing 1,067,120 measurement points.

Strain accuracy and precision were calculated as the mean absolute error (MAER) and standard deviation of error (SDER) as described by Liu and Morgan ^87^

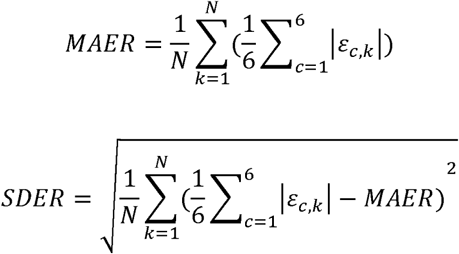

And analogously for the displacements, where N represents the number of points, k is the point in the point cloud, c is the component of strain, and ε is the strain value. Displacement MAER was 10 nm and SDER was 9 nm, strain MAER was 0.036 % and SDER was 0.035 % (full data shown in Figure S4a). The objective minimum values (resulting from minimisation of the ZNSSD function^38^) from DVC were 2.6 × 10^-4^ ± 3.1 × 10^-4^.

A standard zero strain analysis, where two scans were performed sequentially without applying any load to the sample, was performed to test the overall accuracy and precision of the *in-situ* imaging set-up using one lumbar 3-4 sample. A point cloud was generated for the cranial endplate consisting of 1,150,301 points and used for DVC. This gave a displacement accuracy of 85 nm, displacement precision of 63 nm, and strain accuracy and precision of 0.09 % (full data shown in Figure S4b). The objective minimum values from DVC were 7.0 × 10^-4^ ± 3.5 × 10^-4^.

The 1^st^ and 3^rd^ eigenvalues of the strain tensor were calculated and used for measurements of tensile and compressive strain respectively. Maximum shear strain was calculated as: 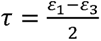, where τ is maximum shear strain, *ε*_1_ is 1^st^ principal strain, and *ε*_3_ is 3^rd^ principal strain. Mean strain values for each endplate are given in Table S3. Mean objective minimum values from DVC are given in Table S4.

Bulk IVD strain was calculated from the change in mean distance between the cranial and caudal VEPs. Initial IVD height was calculated as the distance between the mean (x, y, z) coordinates of the cranial and caudal DVC point clouds. IVD height after each displacement step was calculated by adding the DVC displacements to each point in the point clouds and calculating the new distance between the mean values. The percentage change in height was calculated to give bulk IVD compressive strain, and the values for each sample are given in Table S3.

To determine the direction of strains in the VEP with respect to the loading axis of the joint, the angles between the 1^st^ (tensile) and 3^rd^ (compressive) strain eigenvectors and the z-axis was calculated from 0° - 90°, whereby 0° represents strains acting parallel to the loading axis and 90° strains acting orthogonal to the loading axis. Angles were calculated for each point in the DVC point cloud, and the mean values reported in Results and Table S3.

### Correlative Imaging and Diffraction

Spatially correlated sCT and WAXD were performed at the DIAD beamline, Diamond Light Source^54^ (Figure S12). sCT data was acquired using a pink beam filtered with a Pt strip and 4 mm Al to give a mean beam energy of ∼ 27 keV. An additional 1.4 mm Al and 0.2 mm Cu were used during sample alignment to reduce X-ray dose on the sample. Sample to detector distance was set to 150 mm. Images were collected using scintillator-coupled optics and a PCO.edge 5.5 camera (Excelitas Technologies, Waltham, MA 02451), with a voxel size of 0.54 μm and a field of view of ∼1.38 × 1.16 mm. 2500 projections from 0 to 180° with a 50 ms exposure time were collected per scan. 20 dark and 20 flat field projections were collected at the start and end of each scan for background correction. Images were reconstructed in Savu using filtered back projection, a Fresnel filter, and unsharp mask. Spatial resolution of the images was determined using Fourier shell correlation analysis (Figure S13).

A 17 keV diffraction beam with a 25 μm × 25 μm spot size was used for collection of WAXD data. Data was collected with a PILATUS3X CdTe2M detector ( pixel size 172 μm × 172 μm, Dectris AG, Baden-Daetwill, Switzerland) with an angular position (relative to the sample) covering an azimuthal (χ) range of approximately −67° to 27°, and a sample to detector distance of 32.5 cm, determined through calibration with LaB_6_ ^88^. Diffraction mapping of the endplate was done on a grid of 24 x 17 points with a horizontal step of 54 μm and vertical step of 51 μm, 5 s exposure time was used per point.

### WAXD analysis

WAXD data was processed using moving beam azimuthal integration in the DAWN^89^ (Version 2.33) software package. Azimuthal range was constrained to −67° to 27°, and radial range between 0.79-7.00 Å^-1^ with a binning value of 1000 (including pixel splitting). Resulting one-dimensional datasets of q versus scattering intensity were analysed using custom designed programs^79^ in Python (version 3.10). In each dataset, the (002) peak was isolated by subsampling scattering intensity for q values between 1.75-1.9 Å^-1^. Peaks were modelled by fitting with a Gaussian function using the LMFIT Python library^90^, and only points with fitted peak area greater than 5 (in intensity units) recorded. For these models, (002) d-period was estimated from the model centre q_0_ along q (Å^-1^) by the equation d(002) = 2*π*/q_0_; peak height was estimated as the model amplitude; and peak width was taken as 6*σ* where *σ* is the width term in the Gaussian function. This same method was repeated for the (310) peak, selecting the q range between 2.63-2.89 Å^-1^ and changing the fitted peak area threshold to greater than 30 intensity units (reflecting the higher absolute intensity values of this scattering region in mineralised tissues). For each WAXD map, nanoscale properties were mapped onto two-dimensional images of sCT reconstructions, summed across their x-axis, using the saved motor positions for each WAXD beampath.

Crystallite length and width were calculated from the 002 and 310 peaks respectively, using the following equation:

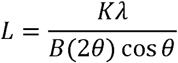

Where *L* is the crystallite size, *K* is a dimensionless shape factor (*K* = 0.9), *λ* is the wavelength of the X-ray beam (*λ* = 0.7348 Å), *B* is the peak width, and 2θ is the Bragg angle^91^.

### Regionalised analysis of microstructure, relative TMD, and DVC strain from sCT data

Regions of interest (ROIs) were taken from the sCT datasets for microstructural analysis and quantification of relative TMD. For whole IVD datasets, 200 voxel (325 µm) cubed regions of interest were chosen at 10 anatomical locations in each endplate (see Fig. 3a) across 8 samples, resulting in 160 regions. For correlative imaging and diffraction experiments, ROIs comprising nine diffraction beam paths were used (Fig. 5e), resulting in 125 x 120 µm rectangles projected through the width of the sample. 15-32 regions were taken across 8 samples, resulting in 194 regions for statistical analysis.

Endplate microstructure was binarized using the interactive thresholding module in Avizo, and ROIs were taken using the extract subvolume module. Greyscale images were masked with the binarized calcified tissue microstructure and mean greyscale value for each ROI was calculated as a representative value for tissue mineral density (TMD). There was significant variation in greyscale values between samples, due to differences in the minimum and maximum intensity of X-rays hitting the detector for each sample, which determines the final image histogram; factors that influence this are sample size and the presence of air bubbles. This meant that inter-sample variation in TMD could not be analysed. Binary images were saved as 3D Tiffs and opened in ImageJ for microstructural quantification using BoneJ (BoneJ2 release 7.0.18) ^92^. Mineralised tissue volume fraction (MV/TV) was calculated using the Area/Volume fraction module. Osteocyte lacunae were removed from the images using a Maximum followed by a Minimum filter (size = 2.5 voxels for I13 sCT data, size = 8 voxels for DIAD sCT data). Mean septal/ trabecular thickness (ST.Th) was calculated using the Thickness module in BoneJ^93,94^. Degree of anisotropy was calculated using the anisotropy module ^95,96^ with 2000 directions, 10000 lines, and a sampling increment of 1.73; anisotropy was not analysed for DIAD data due to the inherent anisotropy in the shape of the ROIs analysed. Purify was run on the regions to ensure the presence of a single connected structure before running the connectivity (Modern) plugin^97,98^ to calculate the connectivity of the ROIs, this was then divided by mineralised tissue volume to give connectivity to mineralised tissue volume ratio (Con/MV). Mean values of DVC measured strain were calculated for each ROI in the whole IVD datasets. Tensile strain (Tension) was calculated as the mean 1^st^ principal strain, compressive strain (Compression) was calculated as the magnitude of mean 3^rd^ principal strain, and shear strain (Shear) was calculated as the mean maximum shear strain in the region.

### Statistical analysis

Statistical analysis was applied to the mean values from regions of interest for each dataset - 160 regions from *in situ* sCT imaging data and 194 regions from correlative imaging and diffraction data. Repeated measures correlation was performed using the rmcorrShiny app^99^ and used to determine common intra-sample associations without violating assumptions of independence of observations^58^, with data grouped by sample (whole or quarter IVD). Each sample was treated as independent to account for variations in mechanical loading, differences in image histograms and subsequent thresholding of microstructure, and the fact that samples from the same animal were from different spinal levels and therefore anatomically distinct. Whole IVD samples for *in situ* sCT data each had 20 regions (Figure S6), and quarter IVD samples for correlative sCT and WAXD had 15-32 regions depending on sample geometry (Figures S14 and S15). The Holm-Bonferroni method was applied in Microsoft Excel and used to deal with familywise error rates^59^, and target p-value set to 0.05.

## Supplementary Information

Document S1. Tables S1 – S9, Figures S1 – S17

Table S1 – sample information for in situ sCT experiments

Figure S1 – experimental set-up for in situ sCT experiments

Table S2 – mechanical loading data for *in situ* sCT experiments

Figure S2 – the impact of mechanical testing rig open frame on imaging

Figure S3 – Fourier shell correlation analysis for I13-2 imaging data

Figure S4 – results from zero-strain DVC analysis

Table S3 – mean results from DVC analysis for each sample

Table S4 – mean objective minimum values from DVC analysis

Figure S5 – Lower resolution DVC strain maps

Figure S6 – The impact of radiation dose on measured strains

Figure S7 – region of interest placement for *in situ* sCT dataset

Table S5 – statistical results from repeated measures correlation analysis of I13-2 imaging and DVC data

Figures S8 – S11 – repeated measures correlation regression graphs for statistically significant correlations from in situ sCT and DVC data

Table S6 – values used to create the radar plots in Figure 4d

Figure S12 – 3^rd^ principal strain maps from sample 7, related to Figure 4c

Table S7 – sample information for the 8 samples analysed from correlative sCT and WAXD experiments

Figure S13 – experimental set-up for correlative imaging and diffraction experiments

Figure S14 – Fourier shell correlation analysis for DIAD imaging data

Figures S15 – S16 – region of interest placement for correlative imaging and diffraction experiments

Table S8 – statistical results from repeated measures correlation analysis of DIAD correlative imaging and diffraction data

Figures S17 – S19 – repeated measures correlation regression graphs for statistically significant correlations from correlative sCT and WAXD data

Table S9 – values used to create the radar plots in Figure 6c

## ACKNOWLEDGEMENTS

Laboratory space and facilities were provided by the Research Complex at Harwell. We thank S. Marussi for his support with the adaptation of the mechanical testing rig. We acknowledge the University of Manchester at Harwell for providing the mechanical testing rig. We thank M. Sherratt for providing samples. We thank J. Brunet, J. Liu, S. Rahmani, S. Marathe, J. Chen, and Y. Zhou for their help on beamtimes. We acknowledge funding from Engineering and Physical Sciences Research Council (EP/V011235/1 & EP/V011006/1), Medical Research Council (MR/R025673/1, MR/V033506/1), Royal Academy of Engineering (CiET 1819/10), Chan Zuckerberg Initiative (CZIF2021-006424), Diamond Light Source beamtimes (MG29633-1, SM29784-6). A.L.P. acknowledges support from the i4health Centre for Doctoral Training.

## AUTHOR CONTRIBUTIONS

Conceptualisation: A.L.P., C.M.D., H.D., N.J.T., B.K.B., A.A.P., H.S.G. and P.D.L.; Methodology: A.L.P., E.N., C.M.D., A.S., H.D. and H.S.G.; Software: A.L.P., E.N. and B.K.B; Validation: A.L.P and E.N.; Formal Analysis: A.L.P. and E.N.; Investigation: A.L.P., E.N., C.M.D., A.S., H.D. and H.S.G.; Resources: H.D., N.J.T., B.K.B., H.S.G. and P.D.L.; Data Curation: A.L.P.; Writing – Original Draft: A.L.P., E.N., A.S., F.B., B.K.B., A.A.P., H.S.G. and P.D.L.; Writing – Review & Editing: All authors; Visualisation: A.L.P., E.N. and A.S.; Supervision: F.B., H.S.G. and P.D.L.; Project Administration: H.S.G. and P.D.L.; Funding Acquisition: N.J.T., B.K.B., A.A.P., H.S.G., P.D.L.

## DATA AVAILABILITY

Example datasets are available on FigShare (Parmenter et al. (2024). Supplementary Data: Multimodal imaging reveals multiscale mechanical interplay in vertebral endplate microarchitecture during intervertebral disc loading. University College London. Dataset. https://doi.org/10.5522/04/26789212). All data (of considerable size) is available from the corresponding authors upon reasonable request.

