## Supplementary Information for "Multimodal imaging reveals multiscale mechanical interplay in vertebral endplate microarchitecture during intervertebral disc loading"

**I13-2 *in situ* sCT experiments and analysis**

***Sample information***

A total of 14 whole IVD samples, derived from 3 animals, were prepared for *in situ* loading experiments at beamline I13-2, Diamond Light Source. The details of the 8 samples analysed for this study are shown in Table S1. 6 Samples were excluded from analysis for the following reasons:

- Samples 1 and 2 – No force data collected (error with loading protocol), poor sample alignment.
- Sample 4 – Vertebra damaged during sample preparation, so unsuitable for mechanical testing. Sample used for zero strain analysis (double projection method).
- Sample 6 – X-ray shutter did not open during second scan (no image taken).
- Sample 9 – Significant movement artefacts in all images.
- Sample 11 – X-ray shutter didn’t open during fourth scan (no image taken).

**Table S1** - Sample information for whole IVD used for in situ sCT and DVC experiments.

| **Sample ID:** | **3** | **5** | **7** | **8** | **10** | **12** | **13** | **14** |
| --- | --- | --- | --- | --- | --- | --- | --- | --- |
| **Spinal Level:** | **L5 – L6** | **L2 – L3** | **L4 – L5** | **T13 – L1** | **T10 -T11** | **L1 – L2** | **L2 – L3** | **L3 – L4** |
| **Spine:** | **1** | **2** | **2** | **3** | **3** | **3** | **3** | **3** |

***Experimental Set-up***

**A photograph of the experimental set-up used at I13-2 is shown in Figure S1. The synchrotron beam energy was** 3 GeV with a current of 300 mA. The undulator insertion device gap was set to 5 mm for imaging and increased to 7 mm during sample alignment to reduce additional dose applied to the sample during alignment. A pink beam filtered with Pyr. Graphite 0.28 mm + 1.06 mm, Al 3.2 mm, Fe 0.14 mm resulted in a mean beam energy of 27 keV. A 500 μm LuAG So14 scintillator coupled to 2x objective lens, and a pco edge 5.5 camera were used, with a total magnification of 4x. Sample to detector distance was 500 mm, and source to sample distance was 230 m. An angular step size of 0.1˚ was used for tomographic imaging, resulting in 1800 projections taken over 180˚ in fly-scan mode.

Each experiment lasted ~100 minutes from when the sample was mounted in the mechanical testing rig on the beamline to acquisition of the final image. Sample hydration was maintained during this time by topping up the phosphate buffered saline (PBS) solution in the Kapton tube around the sample (see Figure S1b and c) between scans using a squeeze bottle with narrow nozzle attachment.

The mechanical testing rig was controlled and monitored from the control room during experiments, with force and displacement data recorded every 500 ms. The peak and relaxed forces recorded during each load/ displacement controlled step for each experiment are provided in Table S2.

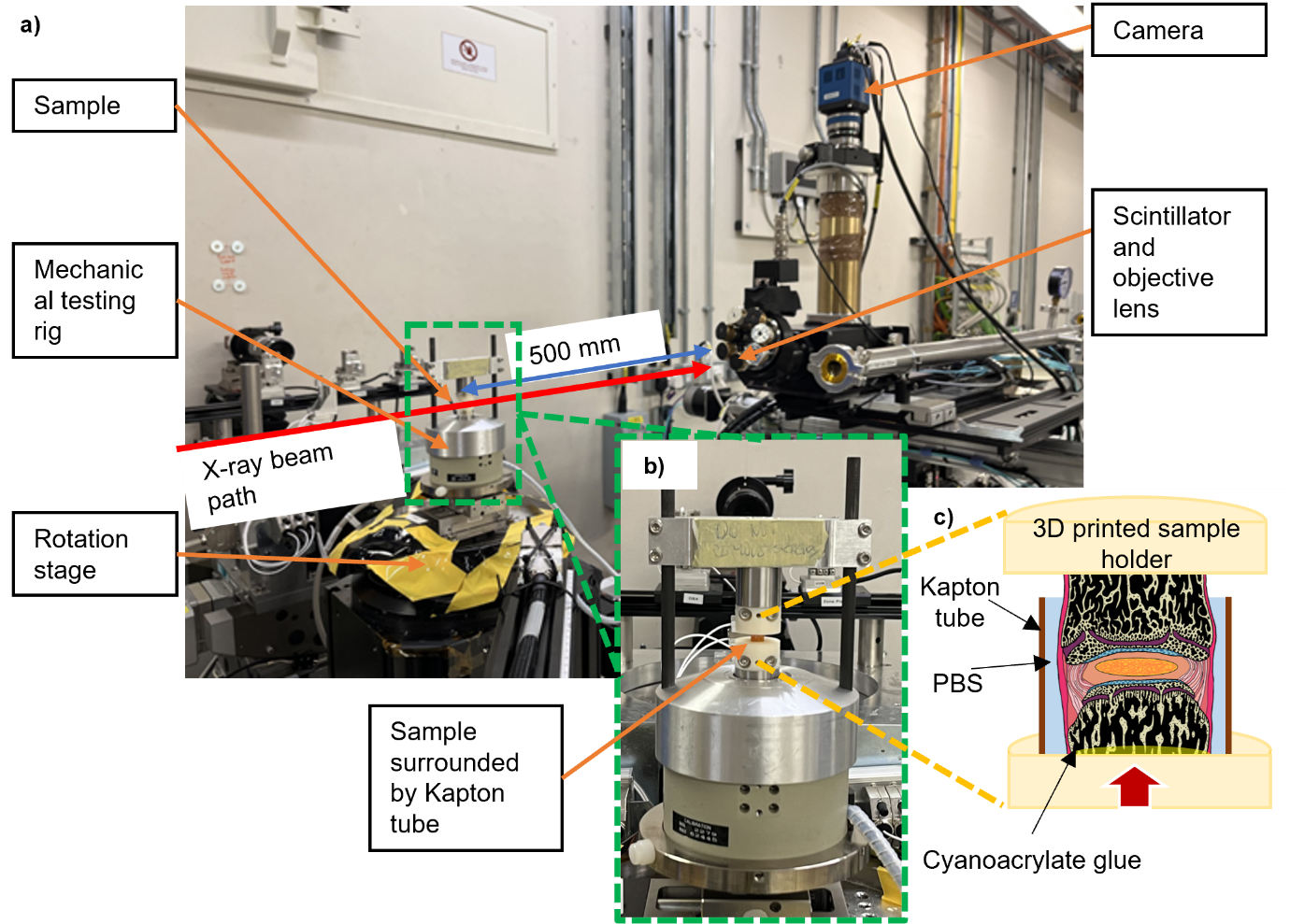

**Figure S1** – **a)** Annotated photograph of the experimental set-up at I13-2; **b)** close-up photo of sample environment; c) illustration of sample environment.

***Mechanical testing data***

**Table S2** - Peak and relaxed forces measured during preload and subsequent 40 µm displacement steps for each experiment used in this study.

| **Sample:** | **Preload peak force (N)** | **Preload relaxed force (N)** | **Step 1 peak force (N)** | **Step 1 relaxed force (N)** | **Step 2 peak force (N)** | **Step 2 relaxed force (N)** | **Step 3 peak force (N)** | **Step 3 relaxed force (N)** |
| --- | --- | --- | --- | --- | --- | --- | --- | --- |
| **3** | **1.03** | **0.01** | **0.26** | **-0.06** | **0.4** | **-0.05** | **1.47** | **1.14** |
| **5** | **1.05** | **-0.01** | **0.98** | **0.01** | **1.08** | **0.02** | **1.11** | **0.01** |
| **7** | **1.01** | **0.2** | **0.54** | **0.27** | **0.58** | **0.3** | **0.64** | **0.31** |
| **8** | **1.15** | **0.35** | **1.18** | **0.44** | **1.45** | **0.45** | **1.41** | **0.44** |
| **10** | **1.00** | **0.06** | **0.96** | **0.21** | **1.23** | **0.33** | **1.53** | **0.42** |
| **12** | **1.03** | **0.07** | **0.92** | **0.07** | **0.78** | **0.03** | **0.97** | **0.01** |
| **13** | **0.98** | **0.16** | **1.59** | **0.32** | **0.65** | **0.38** | **0.89** | **0.42** |
| **14** | **1.04** | **0.22** | **0.84** | **0.3** | **0.99** | **0.33** | **0.89** | **0.32** |
| ****Mean**** | **1.04** | **0.13** | **0.91** | **0.20** | **0.90** | **0.22** | **1.11** | **0.38** |
| ****Standard deviation**** | **0.05** | **0.11** | **0.37** | **0.16** | **0.33** | **0.18** | **0.30** | **0.33** |

***The impact of carbon pillars on X-ray transmission***

A custom open frame was designed for the Deben CT500 mechanical testing rig used for this experiment, enabling easy access to samples during mounting and hydration top-ups. The open frame pillars used for the experiment were carbon rods with a diameter of 6 mm. The pillars appeared in approximately 18.5 % of tomographic projections, resulting in a reduction in intensity at the detector. When one pillar covers the whole field of view it results in a loss of image intensity of approximately 22 %, calculated from the drop in peak frequency intensity shown in Figure S2. This intensity drop did not cause significant imaging artefacts and so was not corrected for during image reconstruction.

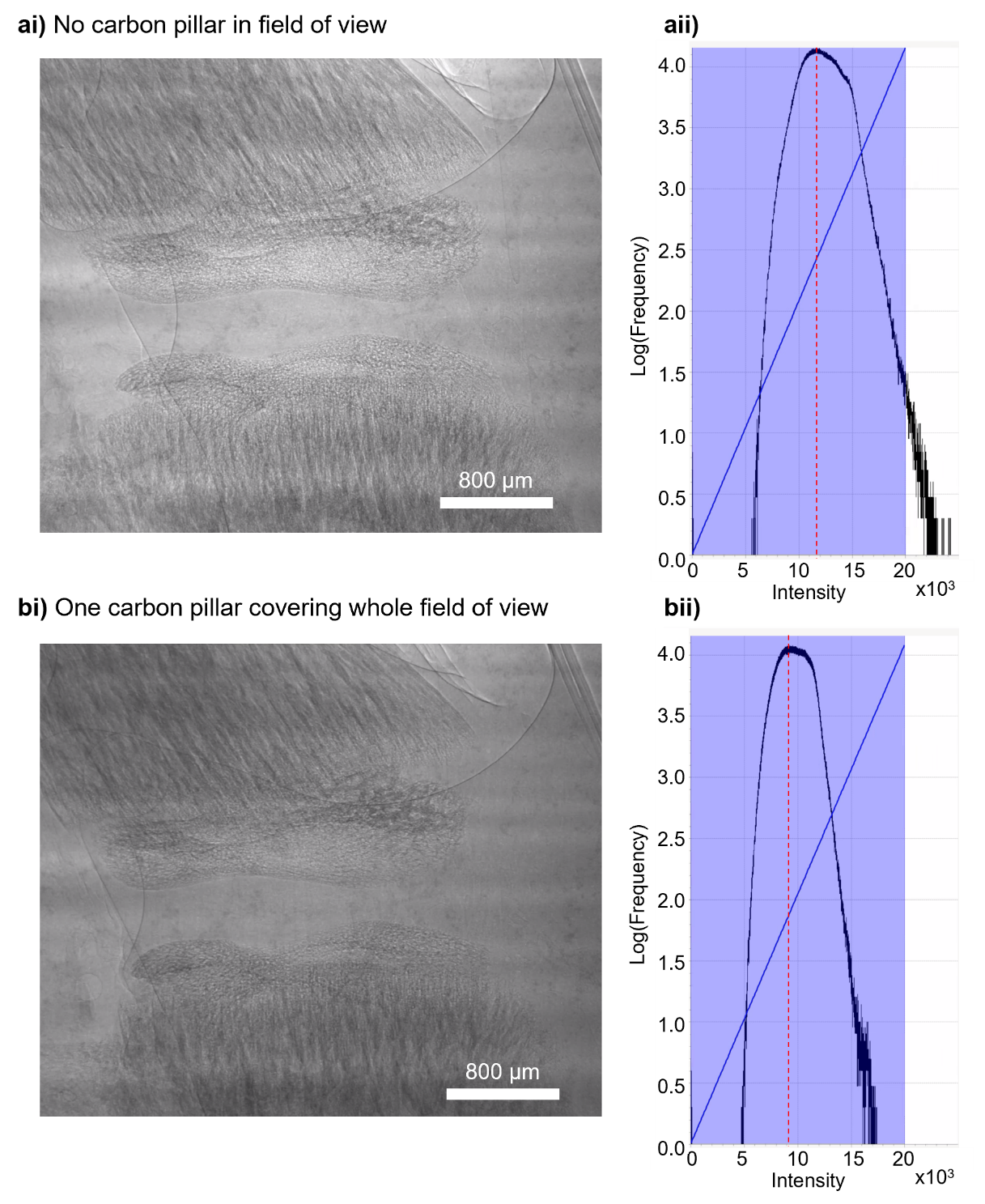

**Figure S2** - **ai)** greyscale intensity projection image and **ii)** histogram of the projection intensity of a sample with no carbon pillar in the field of view, showing a peak frequency intensity of 11800; **bi)** projection image and **ii)** intensity histogram of a projection where one carbon pillar fully covers the field of view, showing a peak frequency intensity of 9200.

***Radiation dose calculation***

Radiation dose for each scan was calculated using the method described in Disney et al 2023^1^. Dose was estimated as

$$Dose \left( Gy \right)=\frac{Exposure time \left( s \right) \times Absorbed energy flux (\frac{J}{sm^{2}})}{Mass per unit area (\frac{kg}{m^{2}})}$$

Total exposure time is given by the number of projections multiplied by 0.1 s exposure time (1801*0.1 = 180.1 s). Using an absorbed energy flux of 7.4x10^-4^, H2O composition, mass per unit area of 1 gcm^3^, and thickness of 5 mm ^1^, total dose for one scan was calculated as 26.6 kG.

***Fourier Shell Correlation analysis***

Fourier shell correlation (FSC) analysis^2^ was performed to estimate the spatial resolution of the imaging datasets. For I13-2 data (voxel size = 1.6 µm), the double projection scan (3600 projections) used for DVC zero strain analysis was split into two sets of 1600 projections (projections containing pillars from the mechanical testing rig were removed) and reconstructed. A 450-voxel cube was taken in the centre of the image (where the Nyquist limit is fulfilled) in the nucleus pulposus (Figure S3a). FSC analysis was performed using image regions containing mineralised tissues, however, the analysis did not converge, likely due to directional noise and imaging artefacts produced by the mineralised tissues. The half bit threshold was used to determine a spatial resolution of 5.5 µm (Figure S3b).

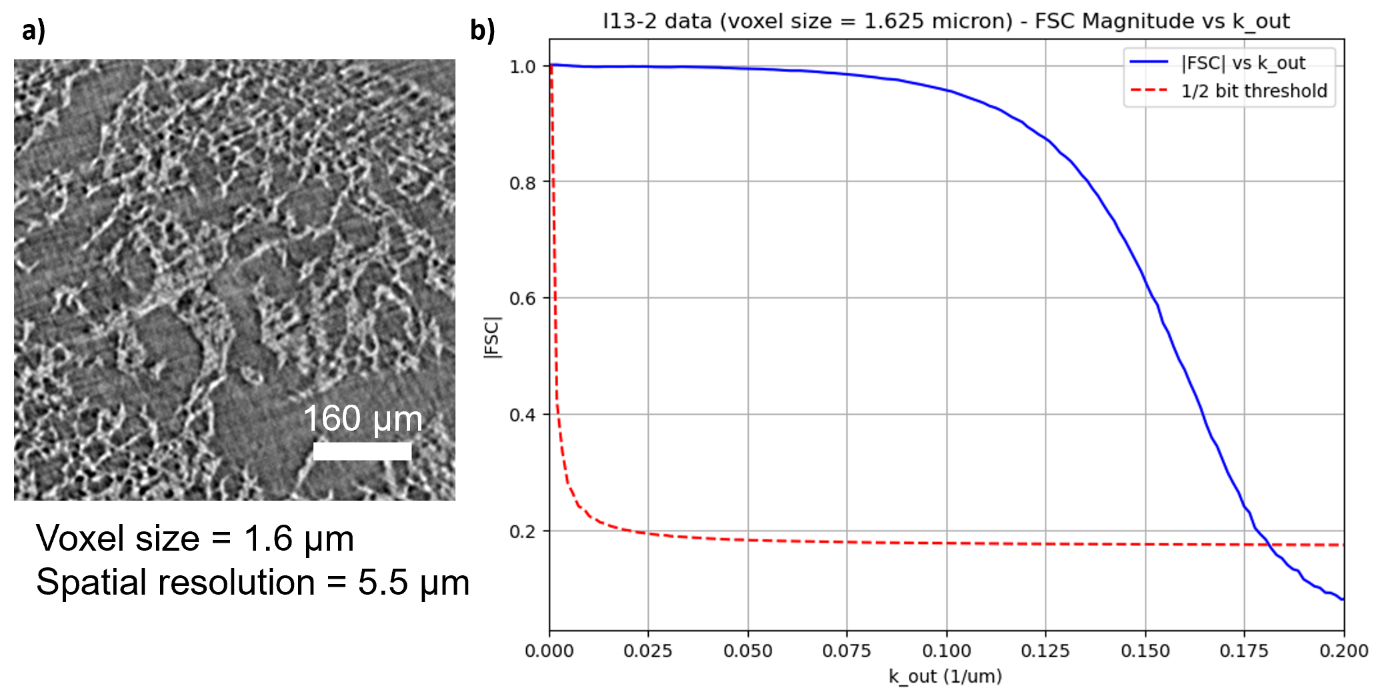

**Figure S3** - **a)** The image region used for FSC analysis, showing microstructures in the nucleus pulposus of the IVD; **b)** magnitude of Fourier shell correlation values versus k values of the shells (blue) with half-bit threshold shown in red – the intersection of the lines corresponds to 1/(spatial resolution).

***Zero strain digital volume correlation analysis***
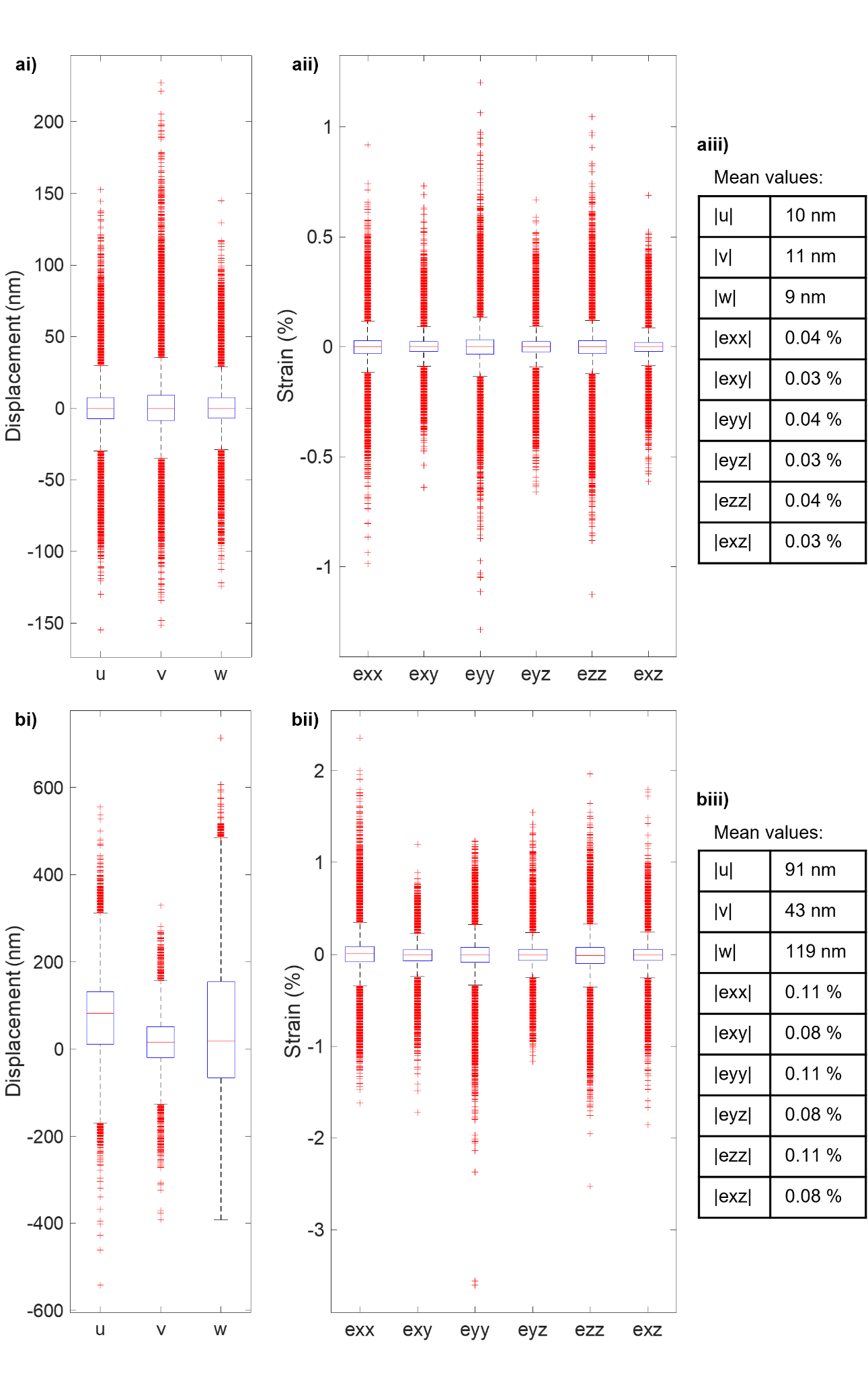

**Figure S4** - Results from **a)** double projection scan zero strain analysis and **b)** repeat scan zero strain analysis showing **i)** box plots of displacements in x, y, and z (u, v, and w respectively), **ii)** boxplots of the 6 strain components and **iii)** mean of the absolute values for each displacement and strain component. Each box plot represents over 1 million data points.

***Digital volume correlation results***

**Table S3** – Initial IVD height, bulk compressive strain in the IVD, mean 1^st^ and 3^rd^ principal strain and the mean angle of their respective eigenvectors with the respect to the loading (z) axis and mean maximum shear strain for the cranial and caudal endplates for each displacement step for the 8 samples analysed.

| Sample ID | Displacement Step | Initial IVD height (µm) | IVD strain (%) | Caudal VEP | | | | | Cranial VEP | | | | |
| --- | --- | --- | --- | --- | --- | --- | --- | --- | --- | --- | --- | --- | --- |
|  |  |  |  | 1^st^ principal strain) | | 3^rd^ principal strain | | Shear strain | 1^st^ principal strain | | 3^rd^ principal strain | | Shear strain |
|  |  |  |  | Mean (%) | Angle (˚ ) | \|Mean\| (%) | Angle (˚ ) | Mean (%) | Mean (%) | Angle (˚ ) | \|Mean\| (%) | Angle (˚ ) | Mean (%) |
| 3 | 1 | 1444 | 2.0 | 0.52 | 60.0 | 0.62 | 53.5 | 0.57 | 0.52 | 59.0 | 0.60 | 53.5 | 0.56 |
|  | 2 | 1415 | 1.8 | 0.38 | 60.0 | 0.48 | 52.3 | 0.43 | 0.45 | 58.5 | 0.54 | 52.3 | 0.49 |
|  | 3 | 1390 | 2.2 | 0.36 | 59.9 | 0.44 | 52.8 | 0.40 | 0.45 | 58.9 | 0.54 | 52.8 | 0.50 |
| 5 | 1 | 1056 | 2.9 | 0.41 | 59.7 | 0.52 | 52.4 | 0.46 | 0.59 | 58.0 | 0.68 | 52.4 | 0.64 |
|  | 2 | 1025 | 2.4 | 0.44 | 58.8 | 0.54 | 52.9 | 0.49 | 0.56 | 58.6 | 0.62 | 52.9 | 0.59 |
|  | 3 | 1001 | 2.3 | 0.48 | 58.6 | 0.59 | 53.4 | 0.53 | 0.61 | 58.1 | 0.70 | 53.4 | 0.65 |
| 7 | 1 | 1256 | 1.6 | 1.61 | 58.5 | 1.77 | 57.4 | 1.69 | 1.87 | 58.3 | 2.11 | 57.4 | 1.99 |
|  | 2 | 1236 | 1.5 | 0.45 | 57.9 | 0.50 | 54.7 | 0.47 | 0.56 | 58.6 | 0.64 | 54.7 | 0.60 |
|  | 3 | 1217 | 1.6 | 0.45 | 58.4 | 0.46 | 54.3 | 0.45 | 0.55 | 58.6 | 0.62 | 54.3 | 0.59 |
| 8 | 1 | 1005 | 3.1 | 0.19 | 58.7 | 0.21 | 47.5 | 0.20 | 0.15 | 58.6 | 0.15 | 47.5 | 0.15 |
|  | 2 | 974 | 2.6 | 0.67 | 58.7 | 0.76 | 53.4 | 0.71 | 0.85 | 59.2 | 0.91 | 53.4 | 0.88 |
|  | 3 | 948 | 2.4 | 0.54 | 58.9 | 0.61 | 52.9 | 0.58 | 0.82 | 59.5 | 0.90 | 52.9 | 0.86 |
| 10 | 1 | 769 | 3.0 | 0.92 | 59.2 | 1.14 | 53.6 | 1.03 | 0.85 | 59.0 | 1.03 | 53.6 | 0.94 |
|  | 2 | 746 | 3.1 | 0.90 | 59.3 | 1.02 | 53.3 | 0.96 | 0.83 | 60.4 | 0.94 | 53.3 | 0.89 |
|  | 3 | 723 | 3.3 | 0.80 | 59.0 | 0.96 | 53.4 | 0.88 | 0.73 | 59.3 | 0.91 | 53.4 | 0.82 |
| 12 | 1 | 1184 | 2.7 | 1.54 | 60.3 | 1.32 | 54.6 | 1.43 | 1.96 | 59.6 | 1.61 | 54.6 | 1.78 |
|  | 2 | 1152 | 2.2 | 0.65 | 60.9 | 0.62 | 52.9 | 0.64 | 0.76 | 60.1 | 0.67 | 52.9 | 0.72 |
|  | 3 | 1127 | 2.6 | 0.37 | 60.8 | 0.43 | 51.7 | 0.40 | 0.43 | 59.9 | 0.44 | 51.7 | 0.43 |
| 13 | 1 | 1221 | 1.7 | 0.56 | 58.7 | 0.64 | 54.7 | 0.60 | 0.57 | 58.7 | 0.70 | 54.7 | 0.64 |
|  | 2 | 1201 | 1.2 | 0.80 | 59.9 | 0.78 | 55.3 | 0.79 | 1.01 | 59.8 | 0.97 | 55.3 | 0.99 |
|  | 3 | 1186 | 1.8 | 0.37 | 58.8 | 0.42 | 53.4 | 0.40 | 0.46 | 58.0 | 0.52 | 53.4 | 0.49 |
| 14 | 1 | 1325 | 2.2 | 0.45 | 59.3 | 0.52 | 53.5 | 0.49 | 0.60 | 58.3 | 0.67 | 53.5 | 0.63 |
|  | 2 | 1296 | 2.0 | 0.44 | 57.6 | 0.49 | 54.7 | 0.46 | 0.59 | 57.5 | 0.65 | 54.7 | 0.62 |
|  | 3 | 1270 | 0.3 | 0.44 | 57.3 | 0.46 | 56.3 | 0.45 | 0.98 | 55.9 | 1.03 | 56.3 | 1.00 |

**Table S4** - Mean objective minimum values output by DVC for the cranial and caudal vertebral endplates for each displacement step.

| **Sample:** | **Mean Objective Minimum** | | | | | |
| --- | --- | --- | --- | --- | --- | --- |
|  | **Step 1 Cranial** | **Step 1 Caudal** | **Step 2 Cranial** | **Step 2 Caudal** | **Step 3 Cranial** | **Step 3 Caudal** |
| **3** | **0.0037** | **0.0040** | **0.0032** | **0.0026** | **0.0035** | **0.0023** |
| **5** | **0.0054** | **0.0036** | **0.0047** | **0.0041** | **0.0056** | **0.0046** |
| **7** | **0.0229** | **0.0239** | **0.0046** | **0.0036** | **0.0046** | **0.0033** |
| **8** | **0.0130** | **0.0137** | **0.0096** | **0.0082** | **0.0094** | **0.0057** |
| **10** | **0.0115** | **0.0158** | **0.0118** | **0.015** | **0.0102** | **0.0132** |
| **12** | **0.0177** | **0.0193** | **0.0047** | **0.0052** | **0.0021** | **0.0026** |
| **13** | **0.0056** | **0.0053** | **0.0091** | **0.0076** | **0.004** | **0.0027** |
| **14** | **0.0059** | **0.0044** | **0.0053** | **0.004** | **0.011** | **0.004** |

***Lower resolution digital volume correlation***

To see overall patterns in the strain field at the macroscale, the displacement field point cloud was randomly down sampled to 2.5 % the number of points, increasing the distance between points from 4.5 voxels (7.3 µm) to 14.6 voxels (23.7 µm). Strains were calculated by polynomial fitting to the 25 closest points, resulting in a mean strain window radius of 42.7 voxels (69.4 µm), and a virtual strain gauge of 115.4 voxels (187.5 µm). At this lower resolution, the increased strain in the central VEP is clearly visible (Figure S5).

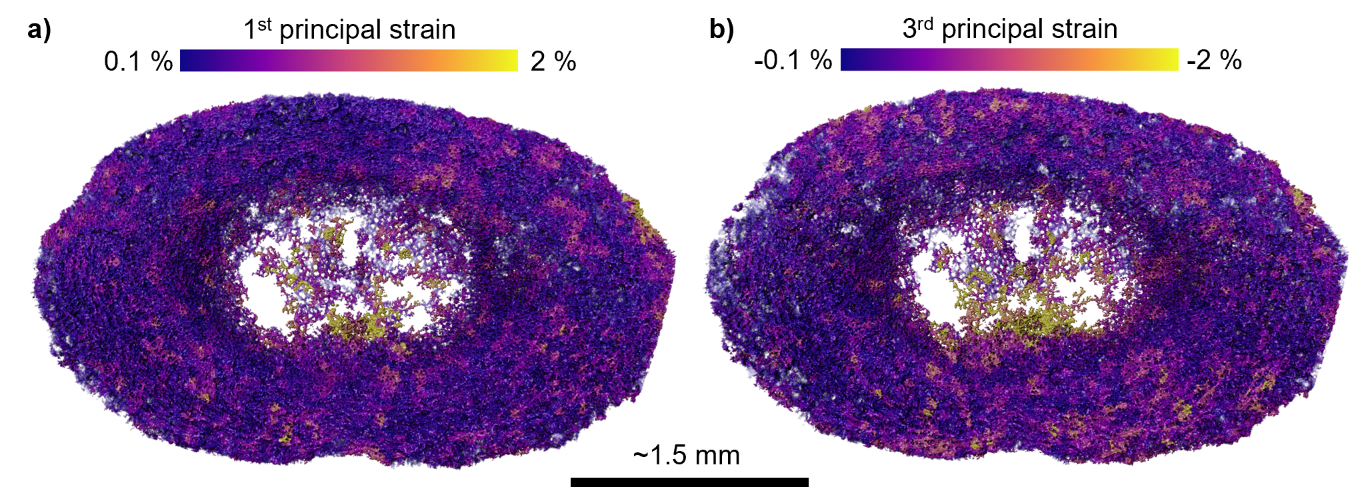

**Figure S5** – Lower resolution **a)** 1^st^ principal strain and **b)** 3^rd^ principal strain in the cranial endplate of sample 5.

***Investigating the impact of radiation dose on digital volume correlation results***

High X-ray dose can cause changes in elastic properties as well as the formation of microcracks in bone^3^. To investigate the impact of radiation dose on measured strain in the VEPs in this study, repeated measures correlation was used to test for statistically significant relationships between mean VEP strains normalised by bulk IVD strain and X-ray dose, and between the percentage of DVC measurement points with a value greater than 10 % strain and X-ray dose (Figure S6). A change in mean strain values generated at each displacement step would indicate a change in elastic properties, and an increase in the number of points with local strains exceeding 10 % would indicate the formation of microcracks^4^.

No statistically significant changes in measured strain values were seen with increasing radiation dose, therefore radiation dose did not have a significant effect on the results obtained in this study.

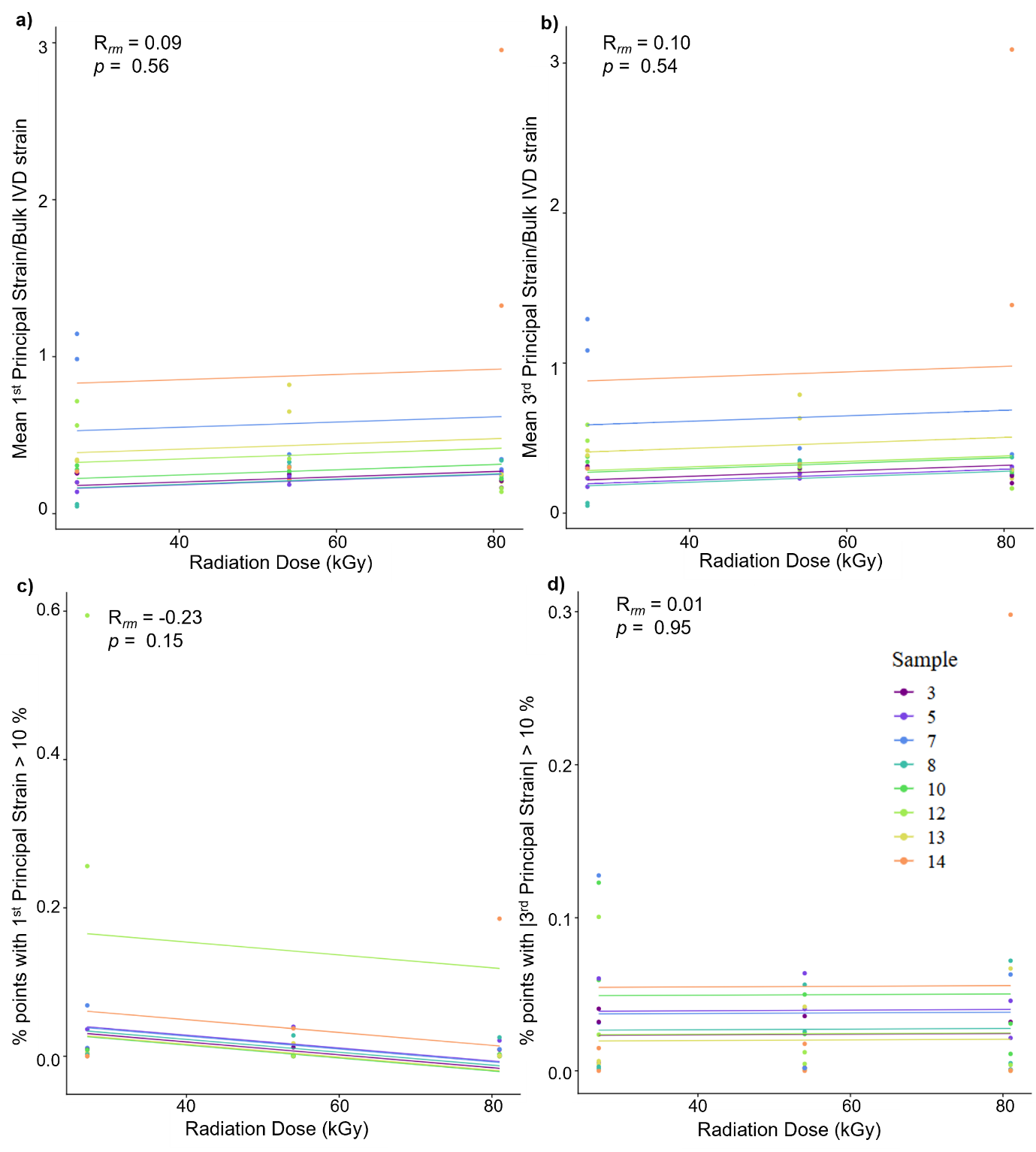

**Figure S6** -Repeated measures correlation regression graphs for **a)** mean 1^st^ principal strain normalised by bulk IVD strain, **b)** mean 3^rd^ principal strain normalised by bulk IVD strain **c)** percentage of DVC measurement points with a 1^st^ principal strain value greater than 10 %, and **d)** percentage of DVC measurement points with a 3^rd^ principal strain magnitude greater than 10 % versus radiation dose. Graphs generated using **the rmcorrShiny app ^5^.**

***Region of interest placement***

**Statistical analysis for the imaging and DVC data collected from I13-2 was performed by taking 10 cubic regions of interest (ROI, side length 325 µm) from each endplate. ROIs were placed in the centre (Figure 3ai 1), anterior inner (Figure 3ai 2), anterior middle (Figure 3ai 3), anterior outer (Figure 3ai 4), lateral inner (Figure 3ai 5), lateral middle (Figure 3ai 6), lateral outer (Figure 3ai 7), posterior inner (Figure 3ai 7), posterior middle (Figure 3ai 9), and posterior outer (Figure 3ai 10) regions. Placement of ROIs was determined by positioning planes along the anterior-posterior and lateral axes of volume renderings of the endplates (Figure S7a) in Avizo** 2022.2 (Thermo Fisher Scientific, Waltham, Massachusetts, U.S.). ROIs were placed equidistantly along these axes, towards the surface between the VEP and the IVD (Figure S7b, c). ROIs were only placed along one lateral side of the VEPs due to larger samples not fitting entirely within the imaging field of view. These larger samples were aligned off centre so that one complete lateral side was capture. Lateral ROIs were taken from the side located the most centrally within the image for all samples.

Each ROI was saved as a 3D binary tif file. The mean greyscale value of the mineralised tissue in each ROI was calculated using the image statistics module in Avizo, with the first 16-bit image of each sample used as input and image statistics calculated within the binary image mask of each ROI. Mean architectural and DVC values were measured for each region (see Methods).

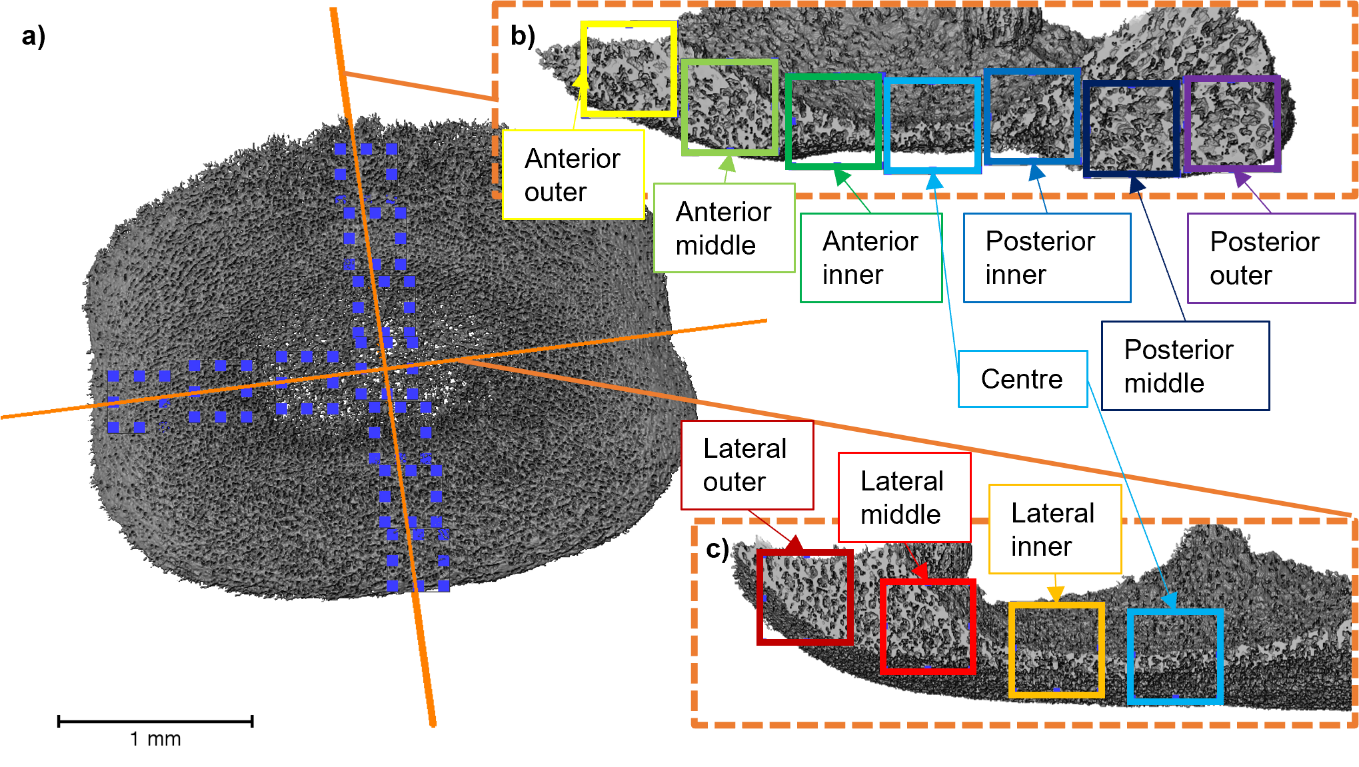

**Figure S7** – Region of interest placement in the caudal VEP of sample 12, **a)** volume rendering of the caudal VEP with orange lines showing the anterior-posterior and lateral axes, ROIs are placed along these axes; **b)** slice through the anterior-posterior axis showing the positions of ROIs along this axis; **c)** slice through the lateral axis showing the positions of ROIs along this axis.

***Statistical analysis***

**The mean values from the 160 VEP ROIs were used for statistical analysis. Statistical analysis was performed using the rmcorrShiny app^5^. ROIs were grouped by whole IVD sample, resulting in 8 groups of 20; each whole IVD sample was treated as independent and common intra-sample relationships were tested using the repeated measures correlation technique^6^. The repeated measures correlation coefficient, 95 % confidence intervals, p-values, rank and alpha level calculated from the Holm-Bonferroni method for each of the 24 relationships tested are given in Table S5. The repeated measures correlation coefficient values were used to generate the heat maps presented in Figures 3 and 4. Regression graphs for statistically significant correlations (p-value less than alpha level) are given in Figures S8-S11, generated using the rmcorrShiny app ^5^.**

**Table S5** – Statistical results from repeated measures correlation analysis of I13-2 imaging and DVC data. 24 relationships were tested between the parameters Location (radial distance from VEP centre), Shear (maximum shear strain), Compression (3^rd^ principal strain magnitude), Tension (1^st^ principal strain), MV/TV (mineralised tissue volume fraction), GSV (image greyscale value), DA (degree of anisotropy), ST.Th (septal/ trabecular thickness), and Con/MV (connectivity to mineralised tissue volume ratio).

| **Parameters** | | **Repeated Measures Correlation Coefficient** | **95 % Confidence Interval** | **p-value** | **Rank** | **Alpha Level (Holm-Bonferroni)**  **(0.05/(24-Rank))** |
| --- | --- | --- | --- | --- | --- | --- |
| Location | MV/TV | 0.76 | 0.678, 0.817 | 1.46e-29 | 1 | 2.17e-3 |
| Shear | GSV | -0.56 | -0.662, -0.443 | 4.11e-14 | 2 | 2.27e-3 |
| Compression | GSV | -0.56 | -0.638, -0.409 | 5.99e-14 | 3 | 2.38e-3 |
| Tension | GSV | -0.53 | -0.659, -0.439 | 1.25e-12 | 4 | 2.50e-3 |
| Location | DA | 0.52 | 0.393, 0.626 | 6.11e-12 | 5 | 2.63e-3 |
| Location | Compression | -0.48 | -0.589, -0.342 | 5.50e-10 | 6 | 2.78e-3 |
| Compression | MV/TV | -0.43 | -0.547, -0.286 | 4.19e-8 | 7 | 2.94e-3 |
| Location | Shear | -0.41 | -0.534, -0.268 | 1.47e-7 | 8 | 3.13e-3 |
| Tension | ST.Th | 0.36 | 0.207, 0.486 | 7.02e-6 | 9 | 3.33e-3 |
| Shear | MV/TV | -0.34 | -0.472, -0.19 | 1.87e-5 | 10 | 3.57e-3 |
| Location | Tension | -0.29 | -0.426, -0.135 | 3.18e-4 | 11 | 3.85e-3 |
| Location | Con/MV | -0.28 | -0.422, -0.129 | 4.10e-4 | 12 | 4.17e-3 |
| Shear | ST.Th | 0.26 | 0.099, 0.397 | 2.00e-3 | 13 | 4.55e-3 |
| Location | ST.Th | 0.23 | 0.071, 0.373 | 5.00e-3 | 14 | 5.00e-3 |
| Tension | MV/TV | -0.19 | -0.34, -0.034 | 0.018 | 15 | 5.56e-3 |
| Compression | ST.Th | 0.17 | 0.01, 0.318 | 0.038 | 16 | 6.25e-3 |
| Location | GSV | 0.16 | 0.002, 0.312 | 0.047 | 17 | 7.14e-3 |
| Compression | Con/MV | 0.13 | -0.026, 0.286 | 0.100 | 18 | 8.33e-3 |
| Shear | Con/MV | 0.10 | -0.063, 0.252 | 0.200 | 19 | 1.00e-2 |
| Tension | DA | 0.10 | -0.061, 0.253 | 0.226 | 20 | 1.25e-2 |
| Tension | Con/MV | 0.04 | -0.12, 0.197 | 0.600 | 21 | 1.67e-2 |
| Compression | DA | -0.03 | -0.186, 0.131 | 0.700 | 22 | 2.50e-2 |
| Shear | DA | 0.03 | -0.134, 0.183 | 0.700 | 23 | 0.05 |

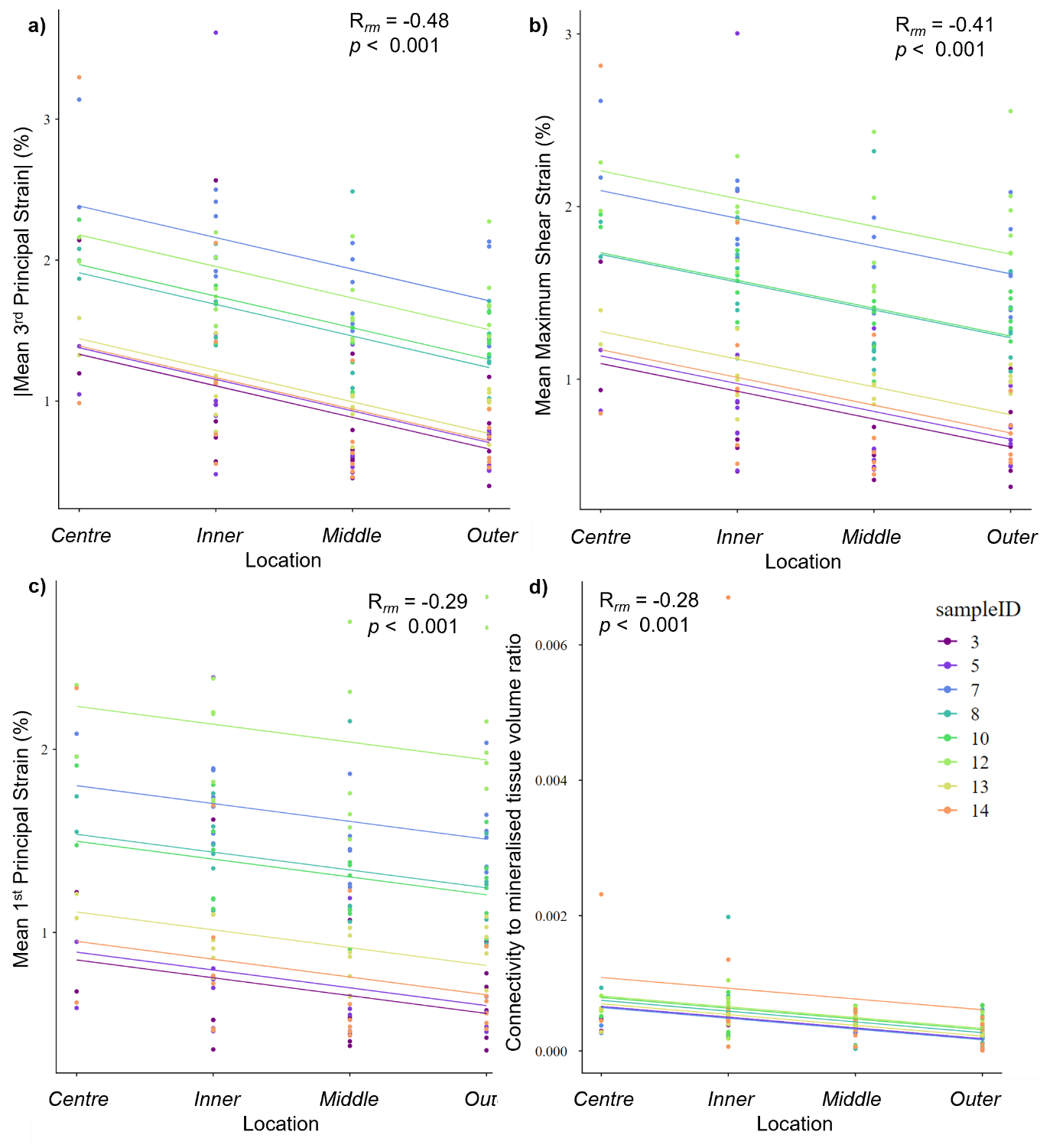

**Figure S8** - Repeated measures correlation regression graphs for **a)** mean compression (3rd Principal Strain), **b)** mean shear (Maximum Shear Strain), **c)** mean tension (1st Principal Strain) and **d)** Con/MV (Connectivity to mineralised tissue volume ratio) vs radial distance from the endplate centre (Location).

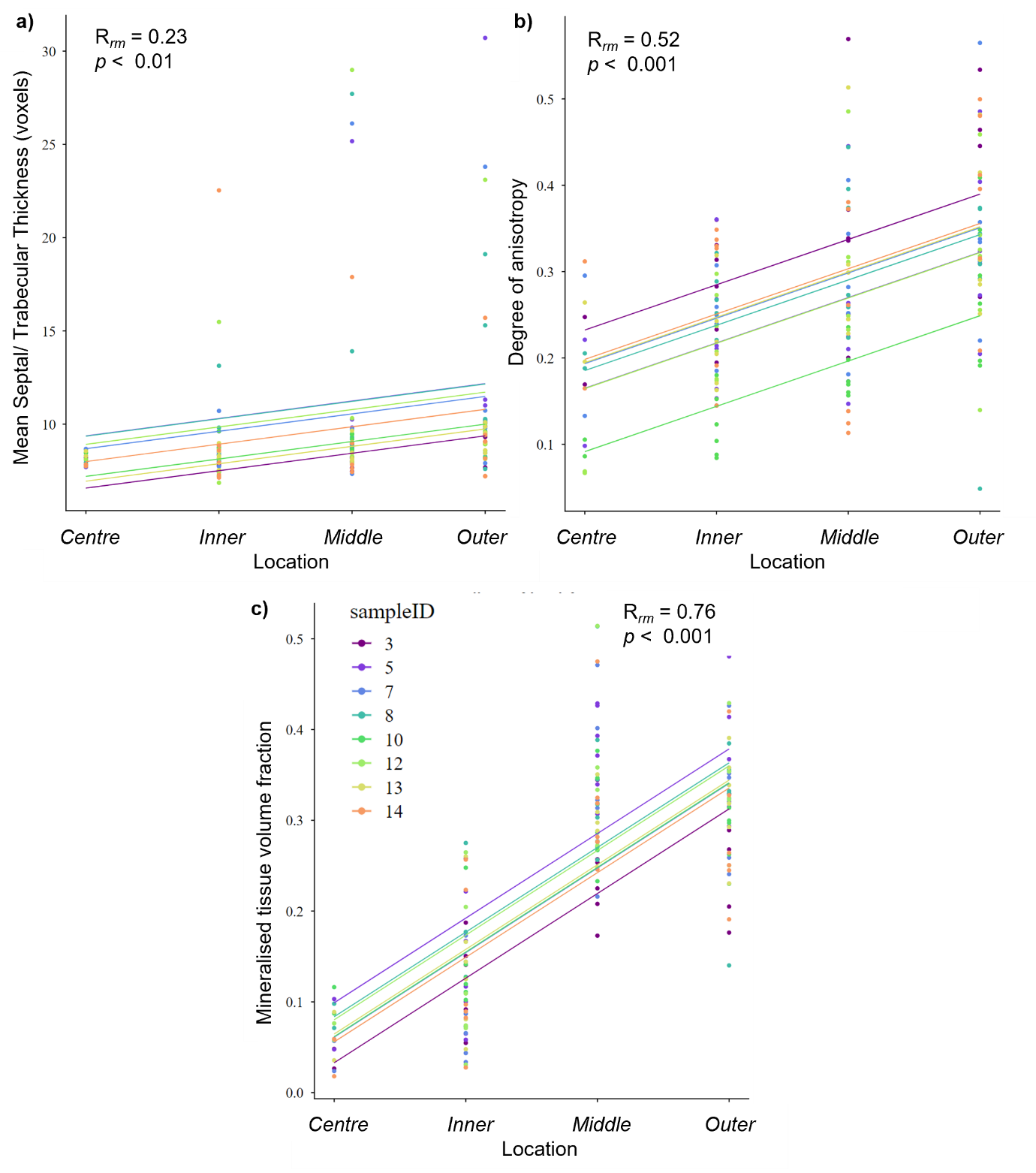

**Figure S9** - Repeated measures correlation regression graphs for **a)** mean ST.Th (Septal/ Trabecular Thickness), **b)** degree of anisotropy, and **c)** MV/TV (Mineralised Tissue Volume Fraction) vs radial distance from the endplate centre (Location).

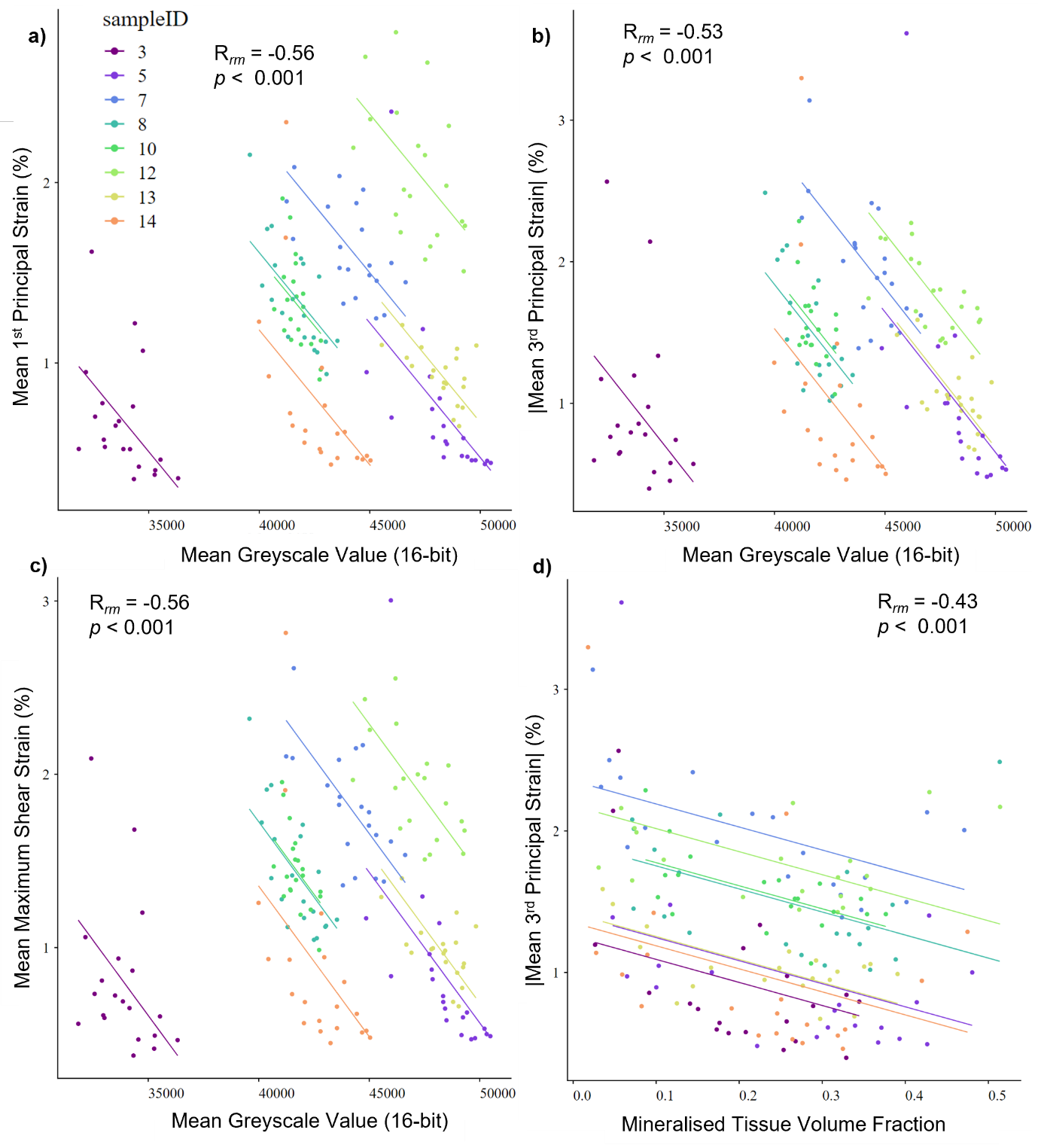

**Figure S10** – Repeated measures correlation regression graphs for **a)** mean tension (1st Principal Strain) vs mean greyscale value **b)** mean compression (3rd Principal Strain) vs mean greyscale value, **c)** mean shear (Maximum Shear Strain) vs mean greyscale value, **c)** mean tension (1st Principal Strain) and **d)** mean compression (3rd Principal Strain) vs MV/TV (Mineralised Tissue Volume Fraction).

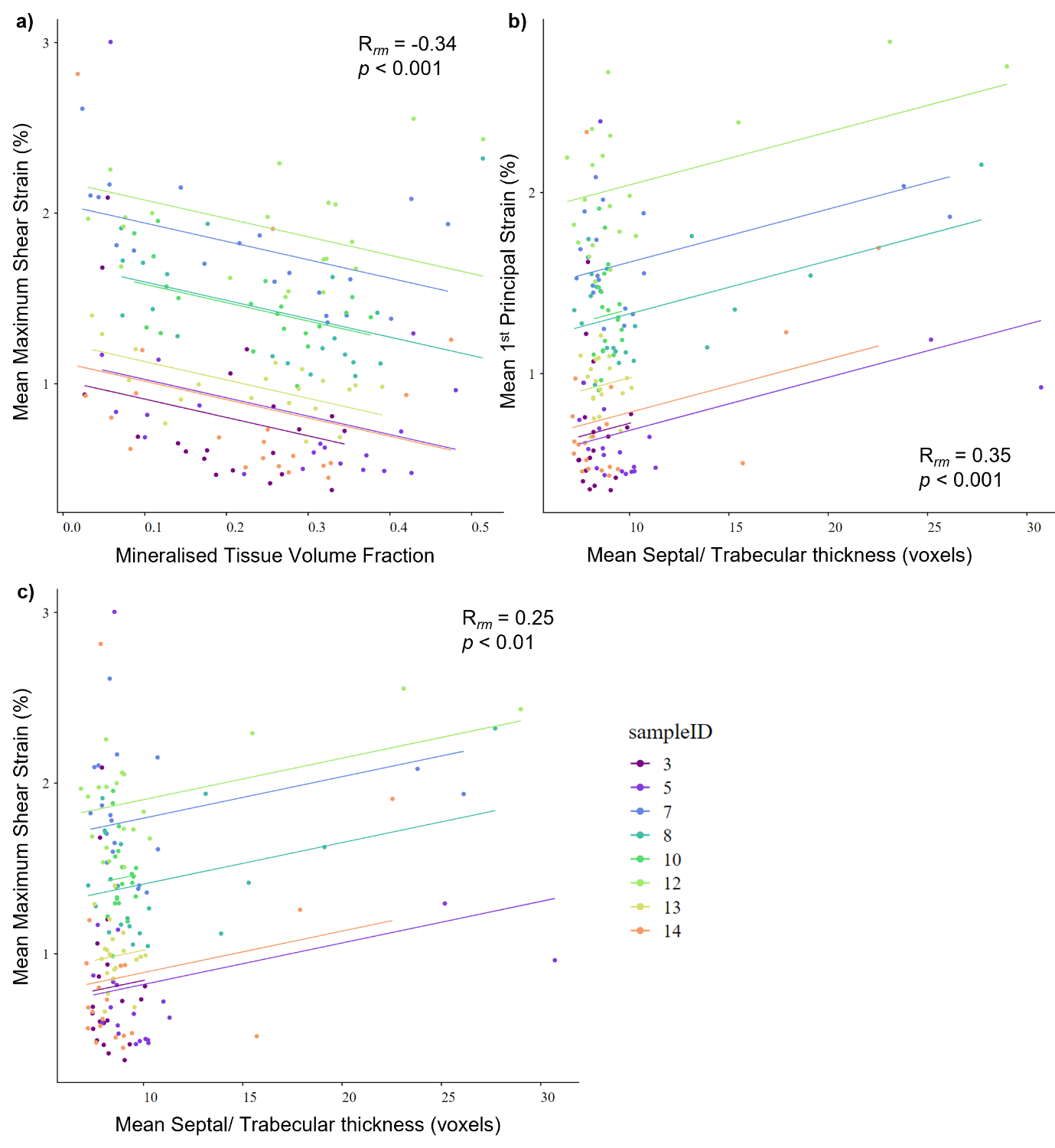

**Figure S11** - Repeated measures correlation regression graphs for **a)** mean shear (Maximum Shear Strain) vs MV/TV (Mineralised Tissue Volume Fraction) **b)** mean tension (1st Principal Strain) vs mean ST.Th (septal/ trabecular thickness), **c)** mean shear (Maximum Shear Strain) vs mean ST.Th (septal/ trabecular thickness).

***Additional data corresponding to Figure 4***

Mean load induced strains after the second displacement step, image greyscale values, mineralised tissue volume, mean septal/trabecular thickness, degree of anisotropy, and connectivity to mineralised tissue volume ratio for the caudal anterior outer, caudal lateral middle, cranial anterior outer, and cranial posterior inner regions of whole IVD Sample 7 are given in Table S6. This data was used to generate the radar plots in Figure 4d. Maps of 3^rd^ principal strain magnitude for these four regions is shown in Figure S12.

**Table S6** – Mean 1^st^ principal strain (Tens.), magnitude of mean 3^rd^ principal strain (Comp.), mean maximum shear strain (Shear), mean image greyscale value (GSV), mineralised tissue volume fraction, mean septal/ trabecular thickness, degree of anisotropy, and connectivity to mineralised tissue volume ration for four regions from Sample 7. These values were used to create the radar plots in Figure 4d.

|  | **Tens. (%)** | **Comp. (%)** | **Shear (%)** | **GSV (16-bit)** | **MV/TV** | **ST.Th (µm)** | **DA** | **Con/MV** |
| --- | --- | --- | --- | --- | --- | --- | --- | --- |
| **Caudal anterior outer** | 1.52 | 1.69 | 1.60 | 44021 | 0.259 | 13.7 | 0.565 | 3.41e-4 |
| **Caudal lateral middle** | 1.45 | 1.60 | 1.53 | 46624 | 0.314 | 13.3 | 0.261 | 6.06e-4 |
| **Cranial anterior outer** | 2.04 | 2.34 | 2.08 | 43630 | 0.426 | 38.7 | 0.312 | 4.27e-5 |
| **Cranial posterior inner** | 1.90 | 2.51 | 2.10 | 41241 | 0.034 | 12.6 | 0.034 | 4.63e-4 |

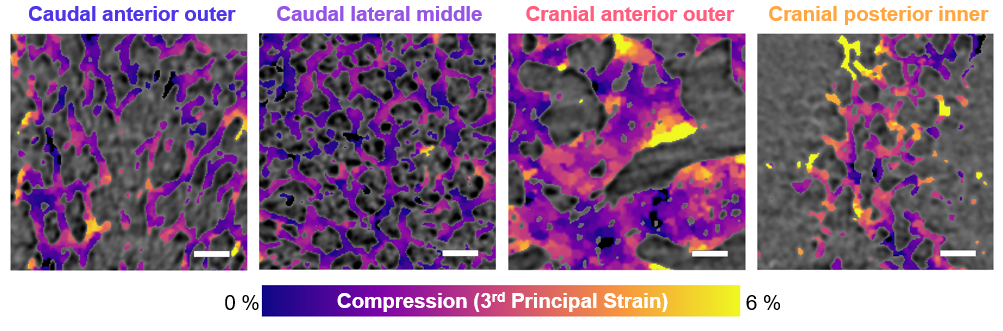

**Figure S12** - 3rd principal strain magnitude after the second displacement step for four regions from Sample 7, corresponding to the 1^st^ principal strain maps shown in Figure 4c.

**DIAD correlative imaging and diffraction experiments and analysis**

***Sample information***

**A total of 26 samples were prepared for correlative sCT and WAXD experiments at DIAD.** The details of the 8 samples analysed for this study are shown in Table S7. 18 samples were excluded from analysis for the following reasons:

- **Sample 1 – Hydration not maintained.**
- **Sample 2 – Glue failed; sample used to test WAXD exposure times.**
- **Sample 3 – Sample too thick, facet joints not fully removed.**
- **Sample 4 – Sample broke when being secured in loading rig.**
- **Sample 6 – Very thin sample broke during loading.**
- **Sample 7 – Significant movement artefacts in sCT images.**
- **Sample 8 – Sample broke when being secured in loading rig.**
- **Sample 9 – Wrong part of sample scanned.**
- **Sample 10 – Sample broke when being secured in loading rig.**
- **Sample 11 – Hydration not maintained, sample broke.**
- **Samples 14 and 15 - Samples broke when being secured in loading rig.**
- **Sample 18 - Sample broke when being secured in loading rig.**
- **Samples 22-26 - Samples broke when being secured in loading rig or glue didn’t hold.**

**Table S7** - Sample information for the 8 samples analysed from correlative sCT and WAXD experiments.

| **Sample ID:** | **5** | **12** | **13** | **16** | **17** | **19** | **20** | **21** |
| --- | --- | --- | --- | --- | --- | --- | --- | --- |
| **Spinal Level:** | **L3-L4** | **L3-L4** | **L1-L2** | **L3-L4** | **L3-L4** | **L1-L2** | **L3-L4** | **L1-L2** |
| **Anterior/ Posterior:** | **Posterior** | **Posterior** | **Posterior** | **Anterior** | **Posterior** | **Anterior** | **Anterior** | **Anterior** |
| **Cranial/ Caudal:** | **Caudal** | **Cranial** | **Cranial** | **Caudal** | **Cranial** | **Cranial** | **Caudal** | **Cranial** |
| **Spine ID:** | **1** | **2** | **2** | **3** | **3** | **3** | **4** | **4** |

***Experimental Set-up***

**A photograph of the experimental set-up used at DIAD is shown in Figure S13. Scan time was 21 minutes for sCT and 35 minutes for a raster map of 17 x 24 WAXD scans, giving a** **total experimental time of ~90 minutes including preload application and stress relaxation time. Sample hydration was maintained by placing a 4 mm diameter Kapton tube around the sample filled with PBS solution (Figure S13b,c).** Sample hydration was maintained during the experiment by topping up the PBS solution between scans using a squeeze bottle with narrow nozzle attachment.

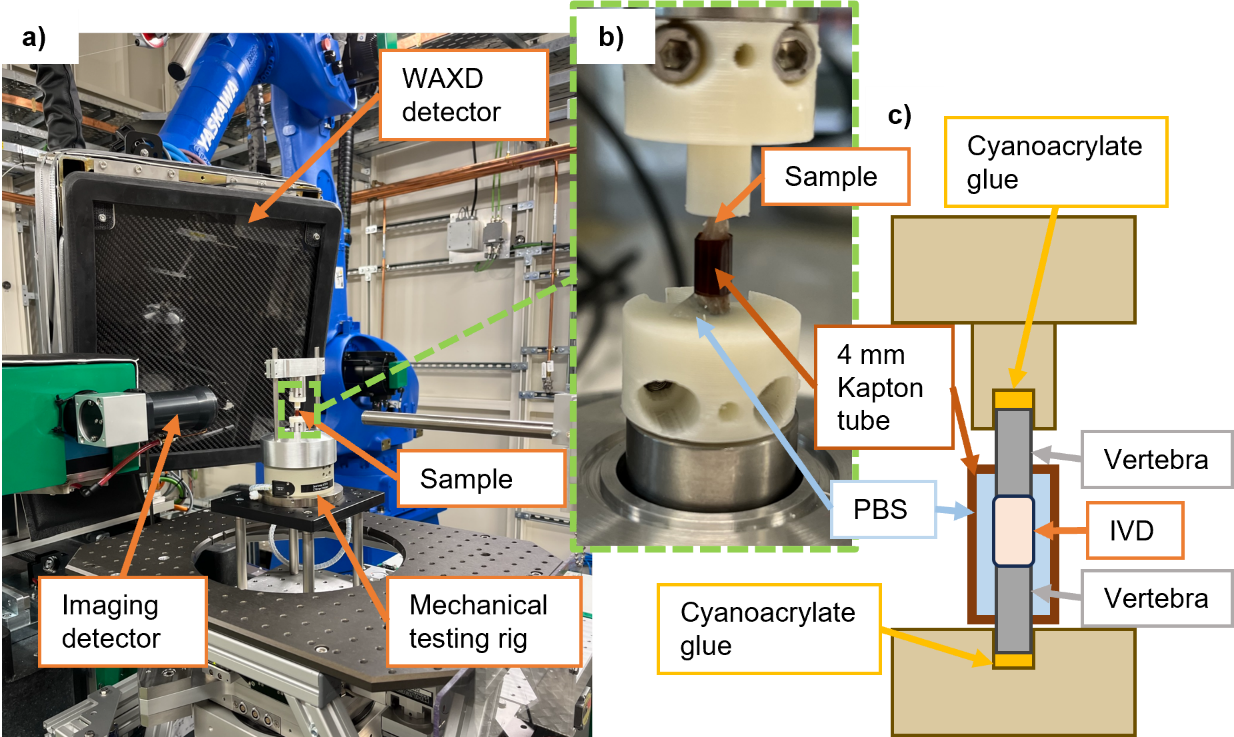

**Figure S13** – Experimental set-up for correlative imaging and diffraction experiments, a) photograph of experimental set-up at DIAD, b) close-up photograph of sample environment, c) illustration of cross-section through sample environment.

***Fourier Shell Correlation analysis***

For FSC analysis of DIAD data (voxel size = 0.54 µm), the scan of sample 5 (2500 projections) was split into two sets of 1150 projections (projections containing pillars from the mechanical testing rig were removed) and reconstructed. A 500-voxel cube was taken in the centre of the image (where the Nyquist limit is fulfilled) in the vertebral bone (Figure S14a). The half bit threshold was used to determine spatial resolution, giving a value of 3.2 µm (Figure S14b). The jaggedness of the FSC curve is likely due to imaging artefacts.
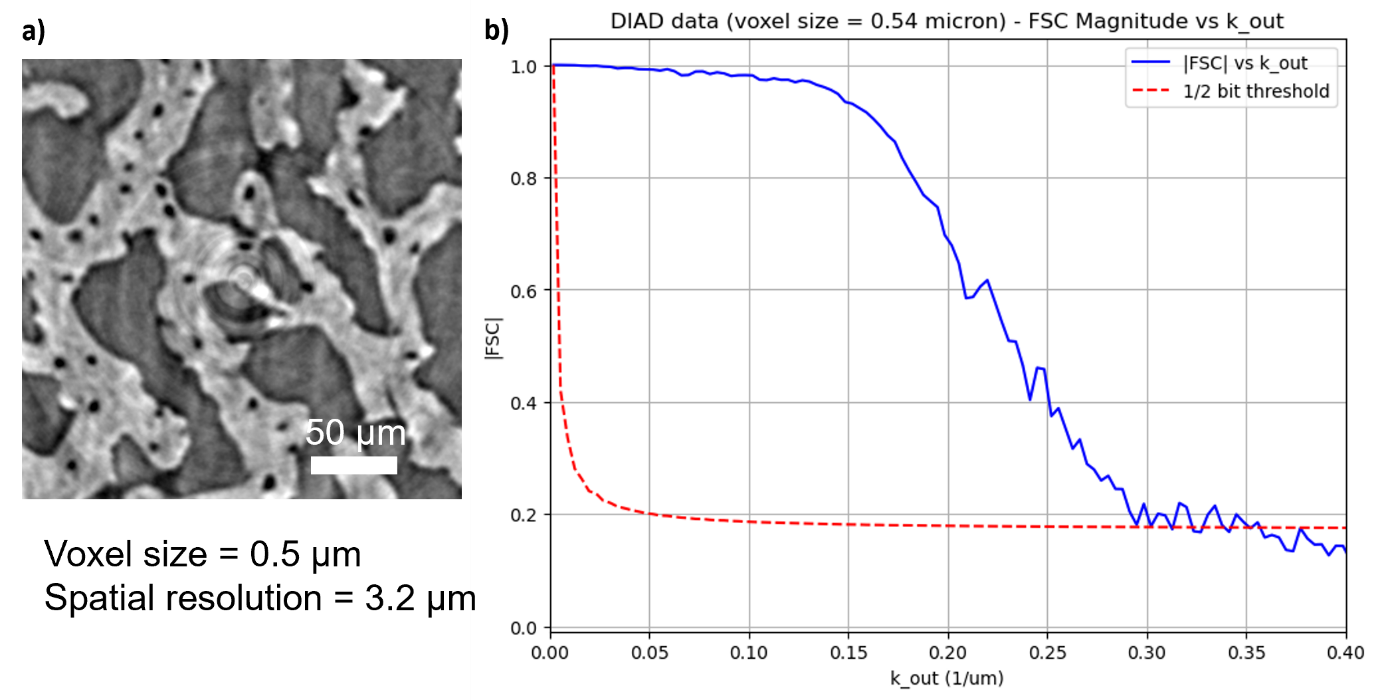

**Figure S14** - a) The image region used for FSC analysis, showing microstructures in the vertebral bone; b) magnitude of Fourier shell correlation values versus k values of the shells (blue) with half-bit threshold shown in red – the first intersection of the lines corresponds to 1/(spatial resolution).

***Region of interest placement***

**Statistical analysis for the imaging and WAXD data collected from DIAD was performed by taking ROIs consisting of 3x3 WAXD beam paths. This resulted in 125 x 120 µm rectangles projected through the width of the quarter IVD sample. 15-32 regions were taken across 8 samples, resulting in 194 regions for statistical analysis. The placement of ROIs for each quarter IVD sample are shown in Figure S15 and S16.**

Each ROI was saved as a 3D binary tif file. The mean greyscale value of the mineralised tissue in each ROI was calculated using the image statistics module in Avizo, with the first 16-bit image of each sample used as input and image statistics calculated within the binary image mask of each ROI. Mean architectural and WAXD values were measured for each region (see Methods). WAXD values corresponding to beam paths that did not pass through the sample but were contained within an ROI were not included when calculating mean values.

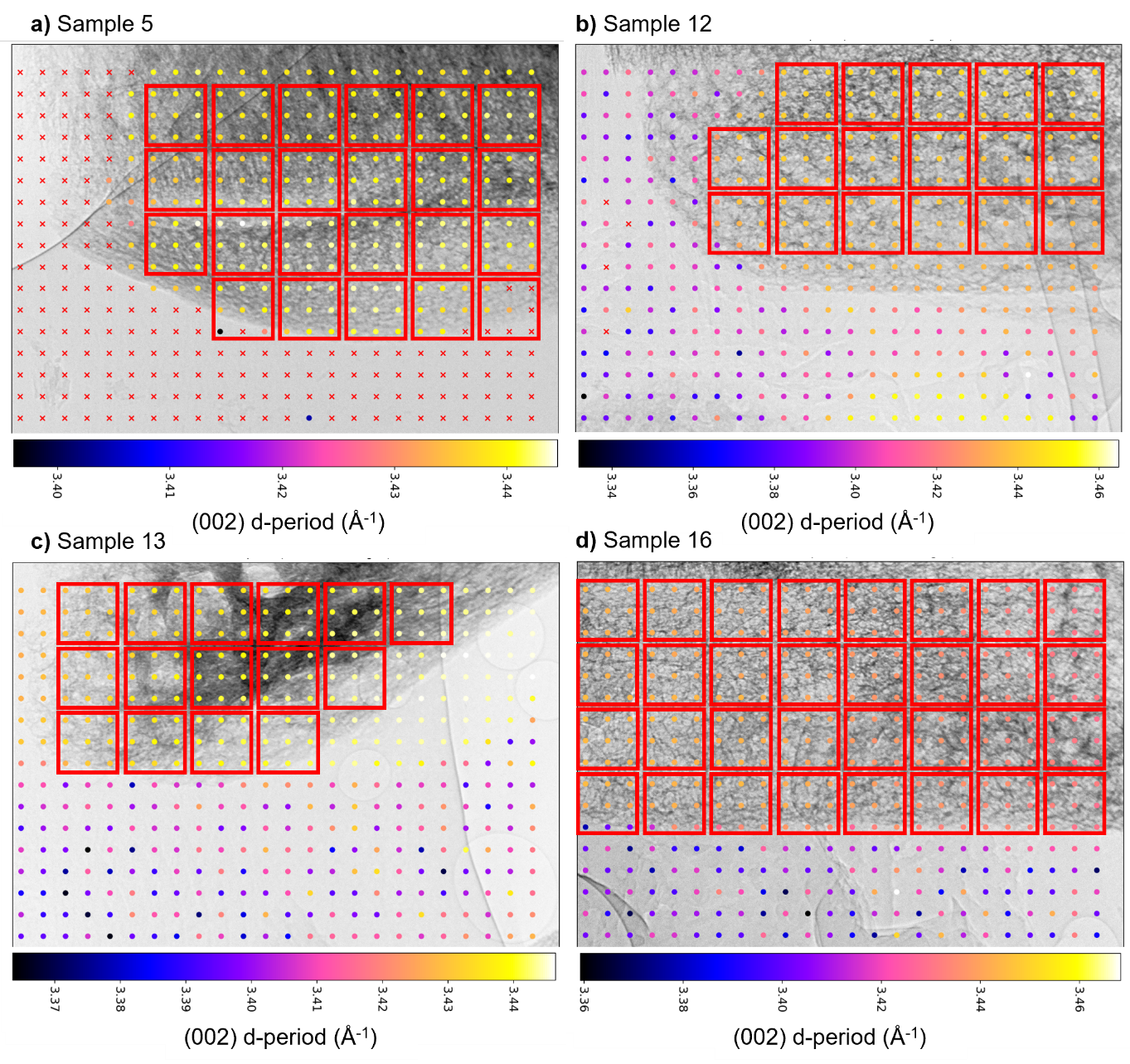

**Figure S15** - Region of interest placement for DIAD data shown on top of radiographs with (002) d-period maps in a) sample 5, b) sample 12, c) sample 13, and d) sample 16.

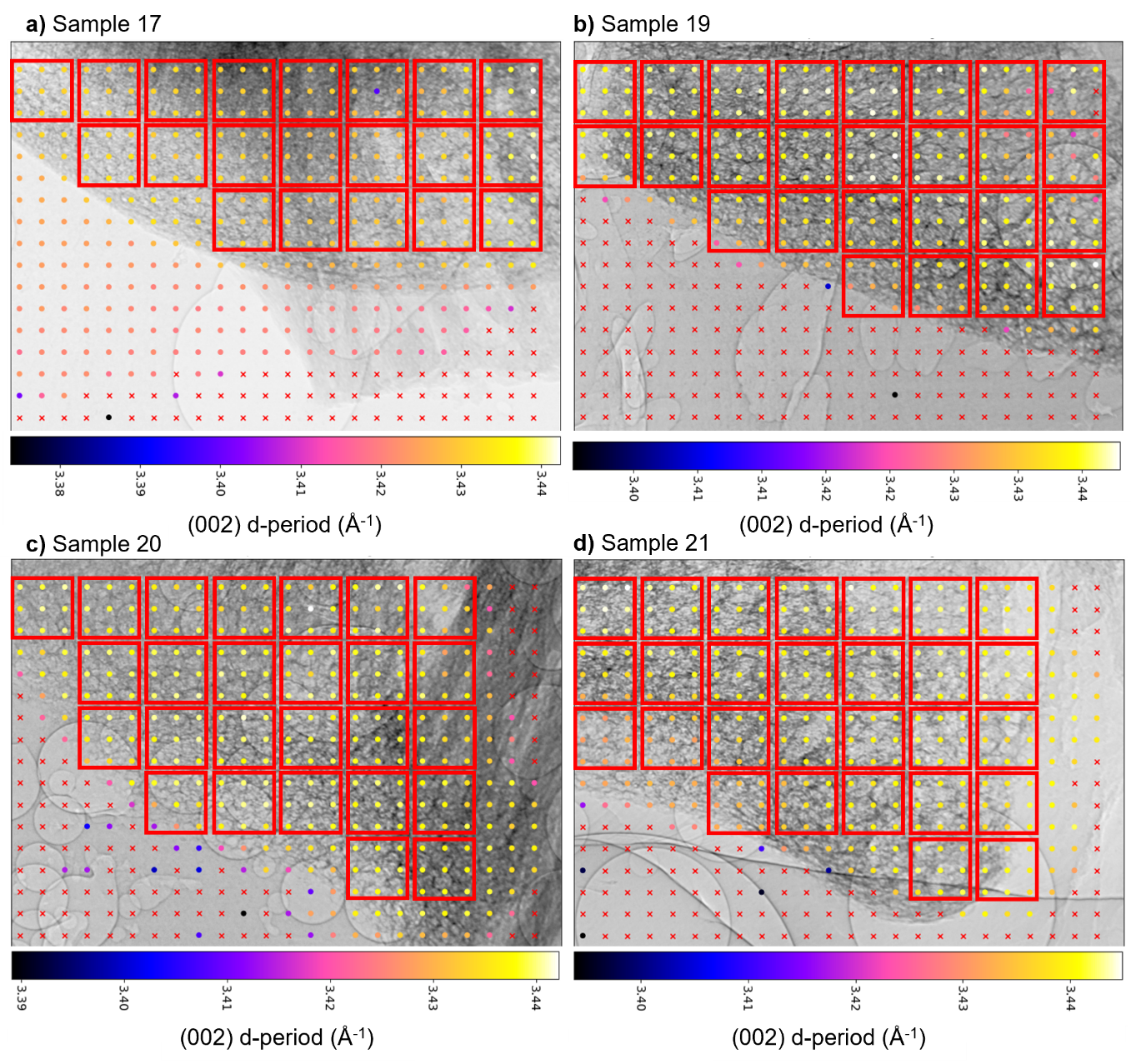

**Figure S16** - Region of interest placement for DIAD data shown on top of (002) d-period maps in a) sample 17, b) sample 19, c) sample 20, and d) sample 21.

***Statistical analysis***

**The mean values from the 194 ROIs were used for statistical analysis. Statistical analysis was performed using the rmcorrShiny app^5^. ROIs were grouped by quarter IVD sample, resulting in 8 groups of 15-32 data points; each quarter IVD sample was treated as independent and common intra-sample relationships were tested using the repeated measures correlation technique^6^. The repeated measures correlation coefficient, 95 % confidence intervals, p-values, rank and alpha level calculated from the Holm-Bonferroni method for each of the 19 relationships tested across the whole dataset are given in Table S8. The repeated measures correlation coefficient values were used to generate the heat map presented in Figure 6a. Regression graphs for statistically significant correlations (p-value less than alpha level) are given in Figures S17-S19, generated using the rmcorrShiny app ^5^.**

**Table S8** – Statistical results from repeated measures correlation analysis of DIAD imaging and WAXD. 19 relationships were tested between the parameters (002) d-period, (310) d-period, Width (crystallite width), Length (crystallite length), MV/TV (mineralised tissue volume fraction), GSV (image greyscale value), ST.Th (septal/ trabecular thickness), and Con/MV (connectivity to mineralised tissue volume ratio).

| **Parameters** | | **Repeated Measures Correlation Coefficient** | **95 % Confidence Interval** | **p-value** | **Rank** | **Alpha Level (Holm-Bonferroni)**  **(0.05/(19-Rank))** |
| --- | --- | --- | --- | --- | --- | --- |
| (002) d-period | Width | 0.44 | 0.317, 0.549 | 2.88e-10 | 1 | 0.0028 |
| (002) d-period | GSV | 0.35 | 0.221, 0.473 | 6.80e-7 | 2 | 0.0029 |
| (002) d-period | Length | 0.34 | 0.207, 0.461 | 1.90e-6 | 3 | 0.0031 |
| Width | GSV | 0.33 | 0.194, 0.45 | 4.58e-6 | 4 | 0.0033 |
| Width | Con/MV | 0.29 | 0.148, 0.412 | 7.44e-5 | 5 | 0.0036 |
| Length | GSV | 0.27 | 0.135, 0.401 | 1.53e-4 | 6 | 0.0038 |
| Width | ST.Th | -0.24 | -0.373, -0.102 | 8.38e-4 | 7 | 0.0042 |
| (002) d-period | ST.Th | -0.24 | -0.368, -0.097 | 0.001 | 8 | 0.0045 |
| (310) d-period | GSV | 0.24 | 0.097, 0.368 | 0.001 | 9 | 0.005 |
| (002) d-period | Con/MV | 0.23 | 0.09, 0.362 | 0.002 | 10 | 0.0056 |
| (002) d-period | MV/TV | -0.21 | -0.34, -0.066 | 0.004 | 11 | 0.0063 |
| Length | MV/TV | 0.18 | 0.035, 0.313 | 0.015 | 12 | 0.0071 |
| Width | MV/TV | -0.18 | -0.312, -0.034 | 0.015 | 13 | 0.0083 |
| (310) d-period | ST.Th | 0.06 | -0.087, 0.199 | 0.4 | 14 | 0.01 |
| (310) d-period | Con/MV | 0.04 | -0.099, 0.187 | 0.5 | 15 | 0.013 |
| (310) d-period | MV/TV | 0.01 | -0.13, 0.157 | 0.8 | 16 | 0.017 |
| Length | Con/MV | 0.01 | -0.138, 0.149 | 0.9 | 17 | 0.025 |
| Length | ST.Th | 0 | -0.147, 0.14 | 1 | 18 | 0.05 |

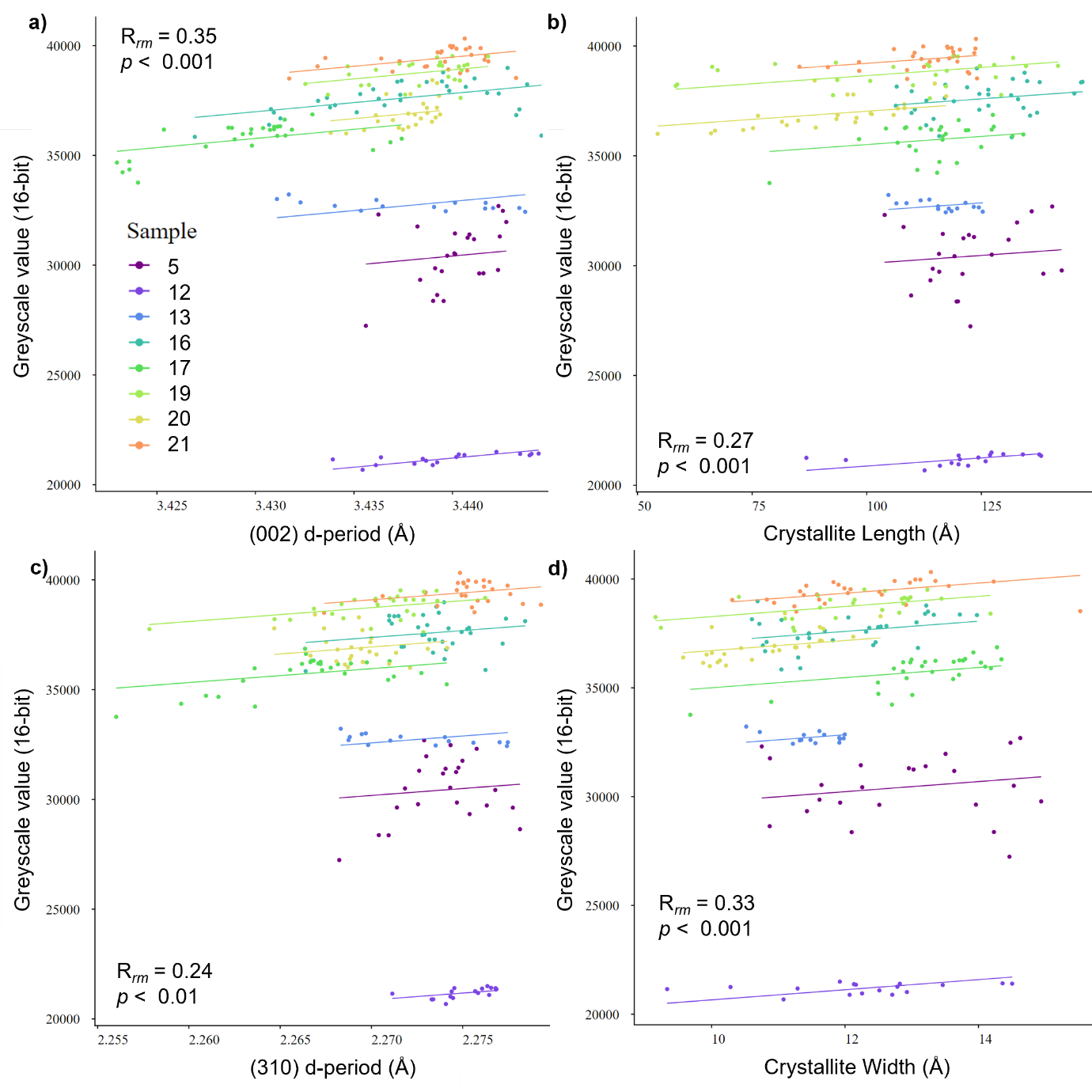

**Figure S17** - Repeated measures correlation regression graphs for greyscale value vs **a)** (002) d-period **b)** crystallite length, **c)** (310) d-period, and **d)** crystallite width.

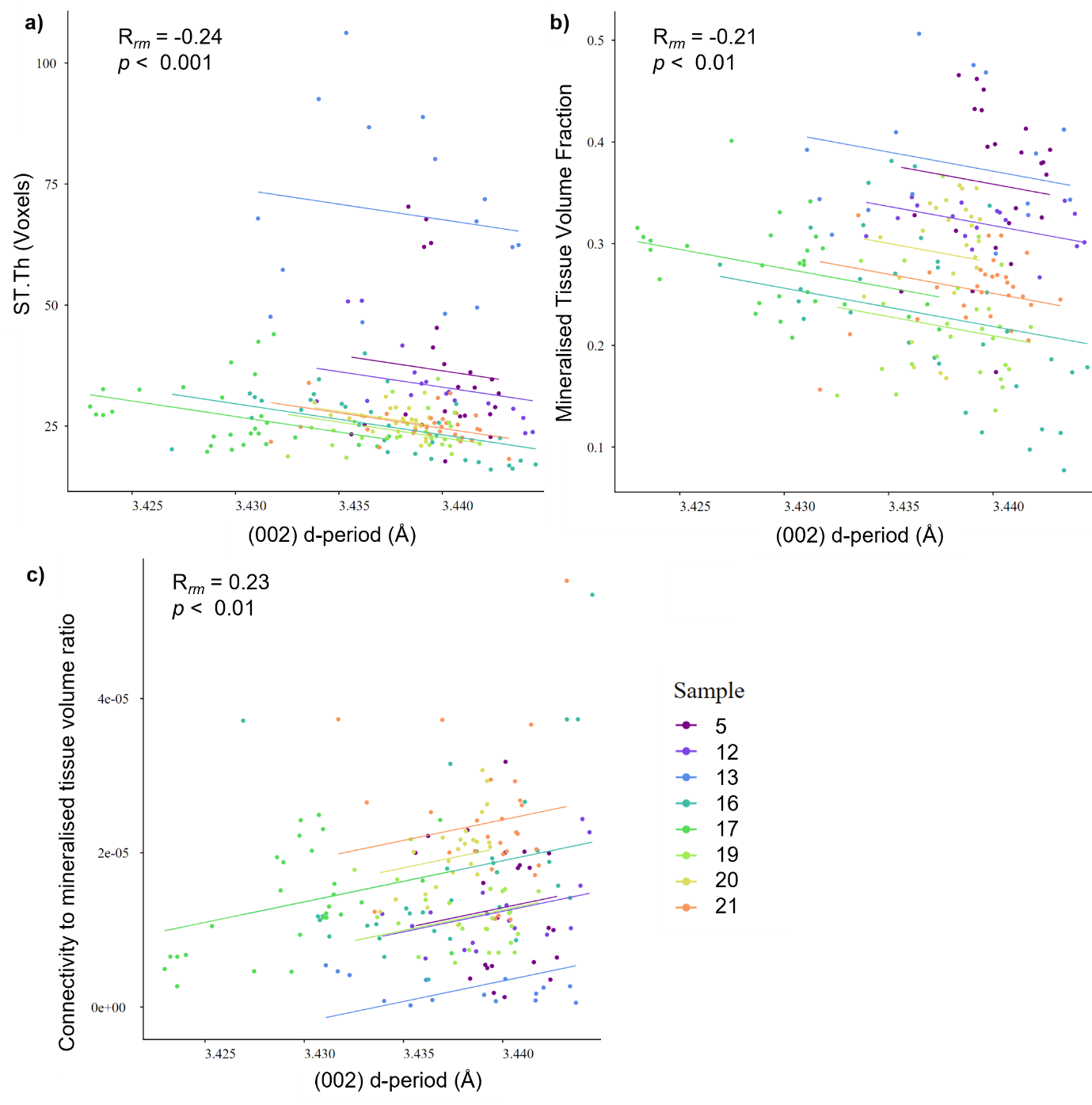

**Figure S18** - Repeated measures correlation regression graphs for **a)** ST.Th (septal/ trabecular thickness), **b)** Mineralised Tissue Volume Fraction, and **c)** connectivity to mineralised tissue volume ratio vs (002) d-period.

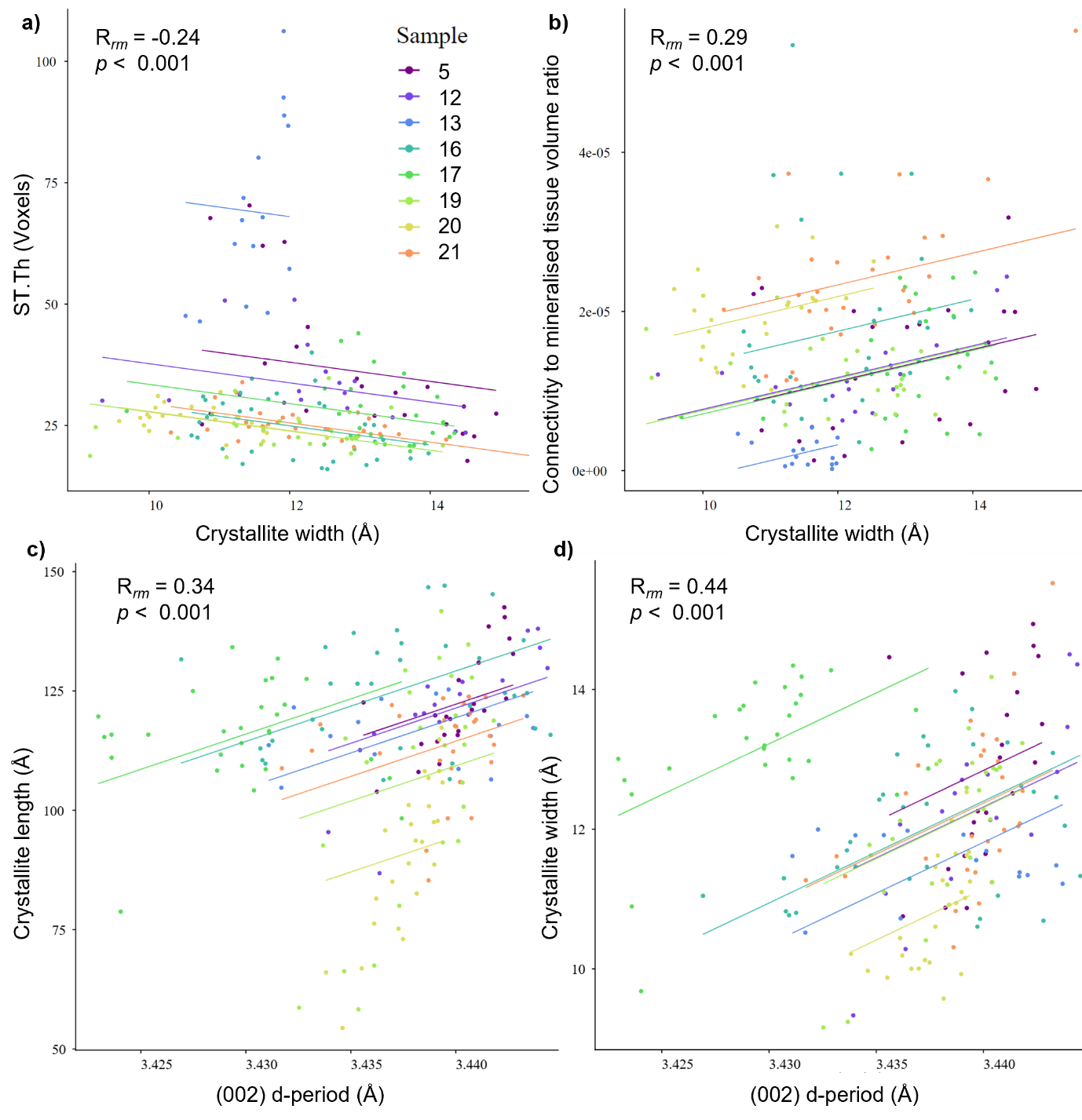

**Figure S19** - Repeated measures correlation regression graphs for **a)** ST.Th (septal/ trabecular thickness) and **b)** connectivity to mineralised tissue volume ratio vs crystallite width; **c)** crystallite length and **d)** crystallite width vs (002) d-period.

***Additional data corresponding to Figure 6***

Mean (002) d-period, (310) d-period, crystallite length and width, greyscale value, mineralised tissue volume fraction, mean septal/ trabecular thickness, and connectivity to mineralised tissue volume ratio for ROI 10, 15, 19, and 30 from sample 16 are given in Table S9. This data was used to generate the radar plots in Figure 6c.

**Table S9** – d-period, crystallite size, and microstructure values used to create the radar plots in Figure 6c.

|  | (002) d-period (Å) | (310) d-period (Å) | Length (Å) | Width (Å) | GSV (16-bit) | MV/TV | ST.Th (µm) | Con/MV |
| --- | --- | --- | --- | --- | --- | --- | --- | --- |
| 17565 ROI 10 (i) | 3.433 | 2.275 | 117 | 11.7 | 37622 | 0.27 | 49 | 1.07e-5 |
| 17565 ROI 15 (ii) | 3.442 | 2.275 | 125 | 12.5 | 37815 | 0.12 | 26 | 1.88e-5 |
| 17565 ROI 19 (iii) | 3.436 | 2.275 | 133 | 12.3 | 37579 | 0.38 | 65 | 3.57e-6 |
| 17565 ROI 30 (iv) | 3.439 | 2.271 | 107 | 10.6 | 38150 | 0.25 | 37 | 1.75e-5 |
